## Supplementary Data for "Pervasive tandem duplications and convergent evolution shape coral genomes"

**Table S1: Statistics of the ONT sequencing data**

| | *Porites lobata* Assembly size of 650Mb | | | *Pocillopora meandrina* Assembly size of 350Mb | | |
|---|---|---|---|---|---|---|
| | Raw ONT reads | Longest reads | Filtlong reads | Raw ONT reads | Longest reads | Filtlong reads |
| Number of runs | 4 MinION + 4 PromethION | | | 4 MinION + 1 PromethION | | |
| Cumulative size | 156 Gb | 21.0 Gb | 21.0 Gb | 62 Gb | 10.5 Gb | 10.5 Gb |
| # of reads | 55,129,048 | 585,173 | 770,592 | 13,260,136 | 191,244 | 286,789 |
| Coverage | 240X | 32X | 32X | 177X | 30X | 30X |
| N50 (bp) | 5,189 | 35,377 | 28,893 | 15,605 | 53,865 | 40,276 |
| Coverage (>50Kb) | 6X | 6X | 4X | 18X | 18X | 10X |

| | *Porites evermanni* Assembly size of 650Mb | | | | |
|---|---|---|---|---|---|
| | Paired-end reads | Mate pairs | | | |
| Sequencer | HiSeq 2500 | HiSeq 4000 | HiSeq 4000 | HiSeq 4000 | HiSeq 4000 |
| Cumulative size | 117.6 Gb | 23.1 Gb | 19.1 Gb | 10.6 Gb | 13.7 Gb |
| # of read pairs | 235,175,203 | 94,673,617 | 77,765,666 | 43,308,999 | 56,684,886 |
| Coverage | 181X | 35X | 29X | 16X | 21X |
| Read size (bp) | 2x250 | 2x150 | 2x150 | 2x150 | 2x150 |
| Insert size (bp) | 400 | 4,000 | 7,000 | 9,000 | 14,000 |

**Table S2: Raw long-read assemblies. A.** ***Porites lobata*****. B.** ***Pocillopora meandrina*****.**

| **A** | SMARTdenovo | | | Flye | | |
|---|---|---|---|---|---|---|
| Subset of reads used | All reads | Filtlong | Longest | All reads | Filtlong | Longest |
| # contigs | 2,225 | 2,149 | 2,098 | 9,845 | 6,771 | 6,287 |
| Cumulative size | 841,373,280 | 776,733,905 | 754,545,205 | 865,703,755 | 870,048,965 | 920,835,732 |
| N50 (L50) | 752,073 (334) | 677,273 (332) | 671,074 (311) | 274,490 (906) | 221,134 (1,143) | 251,501 (1,014) |
| N90 (L90) | 171,297 (1,237) | 144,573 (1,285) | 135,601 (1,287) | 54,588 (3,657) | 72,000 (3,837) | 79,349 (3,575) |
| Max size | 4,191,972 | 3,275,313 | 4,259,348 | 2,045,142 | 1,531,077 | 1,741,775 |
| | Ra | | | RedBean | | |
| Subset of reads used | All reads | Filtlong | Longest | All reads | Filtlong | Longest |
| # contigs | - | 2,992 | 2,693 | 30,253 | 5,792 | 5,429 |
| Cumulative size | - | 374,907,362 | 433,765,815 | 761,471,974 | 668,429,293 | 692,777,752 |
| N50 (L50) | - | 151,282 (792) | 194,050 (681) | 47,890 (3,593) | 383,820 (436) | 393,812 (455) |
| N90 (L90) | - | 68,872 (2,238) | 85,574 (2,032) | 9,095 (18,619) | 51,918 (2,298) | 62,171 (2,154) |
| Max size | - | 897,817 | 1,051,442 | 810,039 | 2,685,953 | 2,863,807 |

| **B** | SMARTdenovo | | | Flye | | |
|---|---|---|---|---|---|---|
| Subset of reads used | All reads | Filtlong | Longest | All reads | Filtlong | Longest |
| # contigs | 388 | 410 | 405 | 2,725 | 2,287 | 2,421 |
| Cumulative size | 371,884,384 | 356,631,128 | 343,867,399 | 399,792,282 | 381,694,599 | 407,450,451 |
| N50 (L50) | 2,617,398 (44) | 2,617,493 (45) | 2,511,303 (44) | 368,823 (282) | 372,078 (256) | 318,427 (340) |
| N90 (L90) | 380,661 (182) | 336,700 (173) | 293,979 (185) | 84,125 (1,216) | 84,025 (1,221) | 90,251 (1,300) |
| Max size | 8,947,962 | 7,928,004 | 6,858,622 | 2,561,294 | 3,845,024 | 2,697,919 |
| | Ra | | | RedBean | | |
| Subset of reads used | All reads | Filtlong | Longest | All reads | Filtlong | Longest |
| # contigs | - | - | - | 12,023 | 1,185 | 1,300 |
| Cumulative size | - | - | - | 382,648,274 | 312,572,668 | 324,887,521 |

| | | | | | | |
|---|---|---|---|---|---|---|
| N50<br>(L50) | - | - | - | 67,474<br>(1,118) | 2,012,258<br>(41) | 1,541,157<br>(60) |
| N90<br>(L90) | - | - | - | 11,305<br>(6,805) | 134,012<br>(259) | 133,570<br>(303) |
| Max size | - | - | - | 1,902,040 | 10,295,455 | 8,452,070 |

**Table S3: Statistics of all assemblies.** The two columns for Haplomerger2 correspond to the two haplotypes generated by the software (a reference haplotype and an alternative haplotype). Two-copy k-mers correspond to duplicated contigs or overlaps in the assembly.

| | *Porites lobata* | | | |
|---|---|---|---|---|
| Method | Raw assembly | Purge Dups | Haplomerger2<br>Reference hap | Haplomerger2<br>Alternative hap |
| # contigs | 2,225 | 1,057 | 1,098 | 1,158 |
| Cumulative size | 851,308,987 | 587,952,820 | 646,152,978 | 588,081,904 |
| N50<br>(L50) | 763,922<br>(333) | 977,395<br>(189) | 2,154,615<br>(84) | 1,959,703<br>(84) |
| N90<br>(L90) | 173,088<br>(1,236) | 302,768<br>(587) | 302,418<br>(357) | 253,346<br>(374) |
| Max size | 4,260,399 | 4,260,399 | 8,615,247 | 7,759,842 |
| BUSCO<br>(N=255) | C:98.4%<br>D:25.1%<br>F:0.4%<br>M:1.2% | C:97.7%<br>D:2.0%<br>F:1.2%<br>M:1.1% | C:98.9%<br>D:2.4%<br>F:0.0%<br>M:1.1% | C:98.5%<br>D:2.0%<br>F:0.4%<br>M:1.1% |
| Number of two-copy k-mers | 77,298,864 | 26,805,171 | 31,535,458 | 28,078,888 |

| | *Pocillopora meandrina* | | | |
|---|---|---|---|---|
| Method | Raw assembly | Purge Dups | Haplomerger2<br>Reference hap | Haplomerger2<br>Alternative hap |
| # contigs | 388 | 388 | 252 | 255 |
| Cumulative size | 376,159,770 | 376,159,770 | 347,233,126 | 340,785,530 |
| N50<br>(L50) | 2,645,240<br>(44) | 2,645,240<br>(44) | 4,753,879<br>(23) | 4,593,186<br>(23) |
| N90<br>(L90) | 392,528<br>(181) | 392,528<br>(181) | 955,142<br>(81) | 905,508<br>(81) |
| Max size | 9,057,404 | 9,057,404 | 11,895,822 | 11,581,782 |
| BUSCO<br>(N=255) | C:98.8%<br>D:7.8%<br>F:0.4%<br>M:0.8% | C:98.8%<br>D:7.8%<br>F:0.4%<br>M:0.8% | C:98.9%<br>D:2.4%<br>F:0.4%<br>M:0.7% | C:98.8%<br>D:2.7%<br>F:0.4%<br>M:0.8% |

| | | | | |
|---|---|---|---|---|
| Number of two-copy k-mers | 25,406,945 | 25,406,945 | 17,008,283 | 16,631,927 |

**Table S4: Statistics of gene prediction on reference and alternative haplotype assemblies.**

| | ***Porites lobata*** | | ***Pocillopora meandrina*** | |
|---|---|---|---|---|
| | *Reference haplotype* | *Alternative haplotype* | *Reference haplotype* | *Alternative haplotype* |
| number of genes | 42,872 | 45,845 | 32,095 | 34,311 |
| number of single-exon genes | 7,496 | 10,906 | 5,341 | 7,135 |
| Average CDS size (bp) | 1,384 | 1,166 | 1,458 | 1,284 |
| Average number of exons per gene | 5.68 | 4.89 | 6.53 | 5.79 |
| BUSCO scores (N=954) | C:95.0% [S:91.3%,D:3.7%] F:1.8% M:3.2% | C:88.7% [S:85.1%,D:3.6%] F:5.8% M:5.5% | C:94.8% [S:92.0%,D:2.8%] F:2.4% M:2.8% | C:91.6% [S:88.6%,D:3.0%] F:4.7% M:3.7% |

**Table S5: BUSCO analyses of raw and haploid assemblies.**

| | *Porites lobata* | | | *Pocillopora meandrina* | | |
|---|---|---|---|---|---|---|
| metazoa_odb10 N=954 genes | Raw assembly | Polished assembly | Haploid assembly | Raw assembly | Polished assembly | Haploid assembly |
| Complete | 92.2% | 95.9% | 95.8% | 89.9% | 95.8% | 95.9% |
| Duplicated | 17.8% | 26.6% | 3.7% | 3.1% | 6.2% | 2.8% |
| Fragmented | 2.9% | 1.6% | 1.5% | 4.7% | 1.9% | 1.8% |
| Missing | 4.9% | 2.5% | 2.7% | 5.4% | 2.3% | 2.3% |

**Table S6: Repeat composition of coral assemblies.**

| | *Porites lobata* | *Pocillopora meandrina* |
|---|---|---|
| bases masked | 56.24 % | 40.30 % |
| Retroelements | 10.02 % | 10.50 % |
| DNA transposons | 17.40 % | 9.00 % |
| Unknown TE | 20.19 % | 13.17 % |
| Simple repeats | 5.83 % | 4.63 % |
| Low complexity | 0.68 % | 1.34 % |
| Satellite | 0.27 % | 0.07 % |
| Rolling-circles | 1.54 % | 0.91 % |
| Small RNA | 0.30 % | 0.67 % |

**Table S7: Data used for comparison of 26 Cnidaria species.** Accession numbers of short-reads datasets used to estimate genome size and abundance of the different OGs in each species, and links to assemblies and annotations downloaded.

| Species | Run accessions (Illumina short-reads) | BioProject accessions (assemblies) | Genome annotations | References |
|---|---|---|---|---|
| *Acropora digitifera* | DRR194164 | | http://spis.reefgenomics.org/download/ | Shinzato, 2011 |
| *Acropora millepora*<br>v1.1<br>*(-> comparative analyses: Figure 1E,F)* | SRR8119879, SRR8119880 | PRJNA473876 | http://coralreefgenomes.jcu.edu.au/ | Ying, 2019 |
| *v2.1*<br>*(-> assembly metrics: Figure 1 B,C,D,G)* | - | PRJNA633778 | https://ftp.ncbi.nlm.nih.gov/genomes/all/GCF/013/753/865/GCF_013753865.1_Amil_v2.1/GCF_013753865.1_Amil_v2.1_translated_cds.faa.gz | Fuller,2020 |
| *Aiptasia sp* | SRR1648348 | | http://aiptasia.reefgenomics.org/ | Baumgarten, 2015 |
| *Amplexidiscus fenestrafer* | SRR5123099, SRR5123102, SRR5123104 | PRJNA354436 | http://corallimorpharia.reefgenomics.org/ | Wang, 2016 |
| *Aurelia aurita* (Atlantic) | - | PRJNA494057 | https://marinegenomics.oist.jp/aurelia_aurita | Khalturin, 2019 |
| *Aurelia aurita* (Pacific) | - | PRJNA494062 | https://marinegenomics.oist.jp/aurelia_aurita_pacific | Khalturin, 2019 |
| *Clytia hemisphaerica* | - | PRJEB30490 | http://marimba.obs-vlfr.fr/ | Leclère, 2019 |
| *Dendronephtya gigantea* | SRR8293698 | PRJNA507923 | GCF_004324835.1 | Jeon, 2019 |
| *Discosoma sp* | SRR5131523, SRR5131524, SRR5131536 | PRJNA354492 | http://corallimorpharia.reefgenomics.org/ | Wang, 2016 |
| *Fungia spp.* | ERR2190368, ERR2190369 | PRJEB23312 | http://ffun.reefgenomics.org/ | Ying, 2018 |
| *Galaxea fascicularis* | ERR2191366 | PRJEB23333 | http://gfas.reefgenomics.org/ | Ying, 2018 |
| *Goniastrea aspera* | ERR2192485, ERR2192486 | PRJEB23371 | http://gasp.reefgenomics.org/ | Ying, 2018 |
| *Hydra vulgaris* | - | | https://research.nhgri.nih.gov/hydra/ | |
| *Montipora capitata* | SRR8497577 | PRJNA509219 | http://cyanophora.rutgers.edu/montipora/ | Shumaker, 2019 |
| *Morbakka virulenta* | - | PRJNA494059 | https://marinegenomics.oist.jp/morbakka_virulenta/ | Khalturin, 2019 |
| *Nematostella vectensis* | - | GCA_000209225.1 | ftp://ftp.ensemblgenomes.org/pub/metazoa/release-43/fasta/nematostella_vectensis | Putnam, 2007 |
| *Orbicella faveolata* | SRR4842281 | PRJNA342412 | ftp://ftp.ncbi.nlm.nih.gov/genomes/Orbicella_faveolata | Prada, 2016 |
| *Pocillopora acuta* | SRR4254617 | | http://ihpe.univ-perp.fr/telechargement/Data_to_downoload.rar | Vidal-Dupiol, 2019 |
| *Pocillopora damicornis* | - | PRJNA454489 | http://pdam.reefgenomics.org/ | Cunning, 2018 |
| ***Pocillopora meandrina*** | | | | **this study** |
| *Pocillopora verrucosa* | SRR11880677 | | http://pver.reefgenomics.org/ | Buitrago-López, 2020 |
| ***Porites evermani*** | | | | **this study** |
| ***Porites lobata*** | | | | **this study** |
| *Porites lutea* | ERR571457, ERR571458, ERR571459 | PRJEB6884 | http://plut.reefgenomics.org/ | Robbins, 2019 |
| *Porites rus* | - | PRJEB23570 | http://gigadb.org/dataset/100462 | Celis, 2018 |
| *Stylophora pistillata* | SRR1980957, SRR1980958 | PRJNA281535 | http://spis.reefgenomics.org/ | Voolstra, 2017 |

**Table S8: Coverage of genome assemblies by syntenic blocks (for contigs with at least 5 genes).**

| | *Porites lobata* | *Porites lutea* | *Pocillopora meandrina* | *Pocillopora verrucosa* |
|---|---|---|---|---|
| *Porites lobata* | - | 397 Mb (78%) | 260 Mb (75%) | 136 Mb (45%) |
| *Porites lutea* | 457 Mb (73%) | - | 239 Mb (69%) | 117 Mb (39%) |
| *Pocillopora meandrina* | 439 Mb (70%) | 305 Mb (60%) | - | 186 Mb (61%) |
| *Pocillopora verrucosa* | 330 Mb (53%) | 216 Mb (42%) | 210 Mb (61%) | - |

**Table S9: Validation of pairs of adjacent duplicate genes and clusters of tandemly duplicated genes.** Two adjacent genes are considered validated if at least one ONT read completely overlaps the two genes. Similarly, a cluster of TDG is considered validated if at least one ONT read completely overlaps all genes of the cluster (Figure S6).

| | | *Porites lobata* | | *Pocillopora meandrina* | |
|---|---|---|---|---|---|
| | | Validated | Not validated | Validated | Not validated |
| Pairs of adjacent duplicated genes | Number | 5,780 (70.3%) | 2,440 (29.7%) | 5,973 (91.2%) | 579 (8.8%) |
| | Average size of the genomic region (bp) | 23,594 | 89,790 | 23,865 | 95,434 |
| | Median size of the genomic region (bp) | 20,781 | 79,530 | 19,990 | 86,723 |
| Clusters of TDG | Number | 1,680 (45.5%) | 2,012 (54.5%) | 1,838 (67.0%) | 903 (33.0%) |
| | Average size of the genomic region (bp) | 27,316 | 144,391 | 31,700 | 150,589 |
| | Median size of the genomic region (bp) | 24,990 | 109,211 | 27,771 | 115,593 |

**Table S10**: Table of calcification-related proteins in non-scleractinian and scleractinian - robust and - complex coral species.

| Hexacorallia | Amt1 | SLC4 | | PMCA | CA | CARP | Neurexin |
|---|---|---|---|---|---|---|---|
| | | β | ɣ | | | | |
| **Corallimorpharia** | | | | | | | |
| *Discosoma sp.* | 1 | 1 | 0 | 2 | 6 | 2 | 4 |
| *Amplexidiscus fenestrafer* | 1 | 1 | 0 | 2 | 6 | 2 | 4 |
| **Actinaria** | | | | | | | |
| *Exaiptasia diaphana* | 2 | 1 | 0 | 3 | 10 | 2 | 3 |
| **Scleractinia - "Robust"** | | | | | | | |
| *Stylophora pistillata* | 5 | 1 | 1 | 3 | 16 | 5 | 3 |
| *Pocillopora verrucosa* | 4 | 1 | 1 | 3 | 14 | 5 | 4 |
| *Pocillopora meandrina* | 5 | 1 | 1 | 3 | 15 | 5 | 3 |
| *Orbicella faveolata* | 4 | 1 | 1 | 3 | 14 | 4 | 4 |
| *Goniastrea aspera* | 4 | 1 | 1 | 3 | 12 | 4 | 3 |
| *Fungia sp.* | 2 | 1 | 1 | 3 | 14 | 4 | 3 |
| **Scleractinia - "Complex"** | | | | | | | |
| *Acropora digitifera* | 2 | 1 | 1 | 3 | 10 | 3 | 3 |
| *Acropora millepora* | 2 | 1 | 1 | 3 | 10 | 3 | 3 |
| *Galaxea fascicularis* | 1 | 1 | 1 | 3 | 9 | 3 | 4 |
| *Montipora capitata* | 2 | 1 | 1 | 3 | 9 | 5 | 3 |
| *Porites lobata* | 3 | 1 | 1 | 3 | 10 | 5 | 5 |
| *Porites lutea* | 3 | 1 | 1 | 1 | 11 | 3 | 4 |

**Figure S1**. K-mer distribution and estimation of genome sizes and heterozygous rates from short reads. (A) *Porites lobata*, (B) *Porites evermanni*, and (C) *Pocillopora meandrina.*

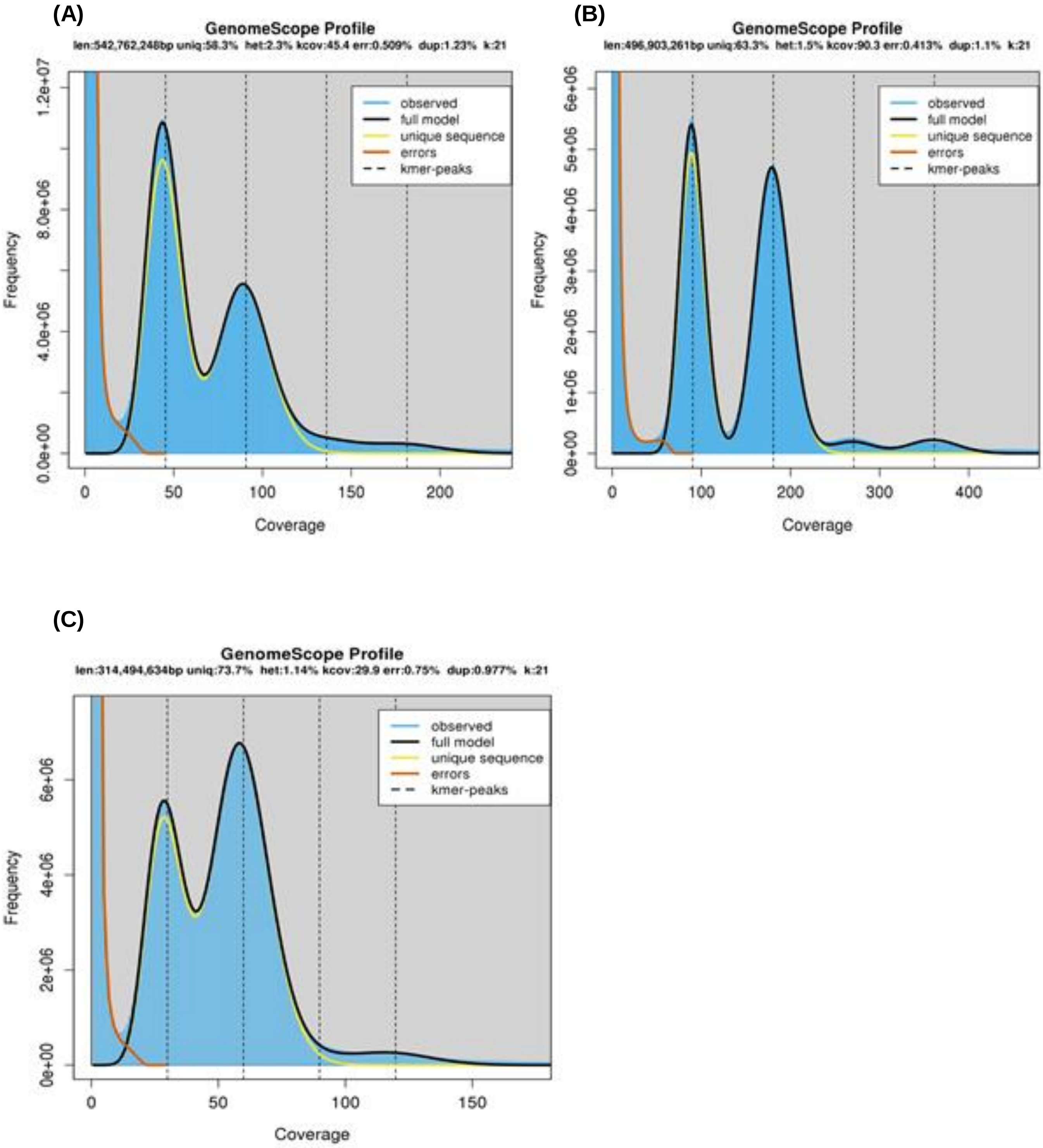

**Figure S2**. KAT plot of *Porites lobata* assemblies. (A) polished nanopore assembly. (B) haploid version of the genome.

**A**

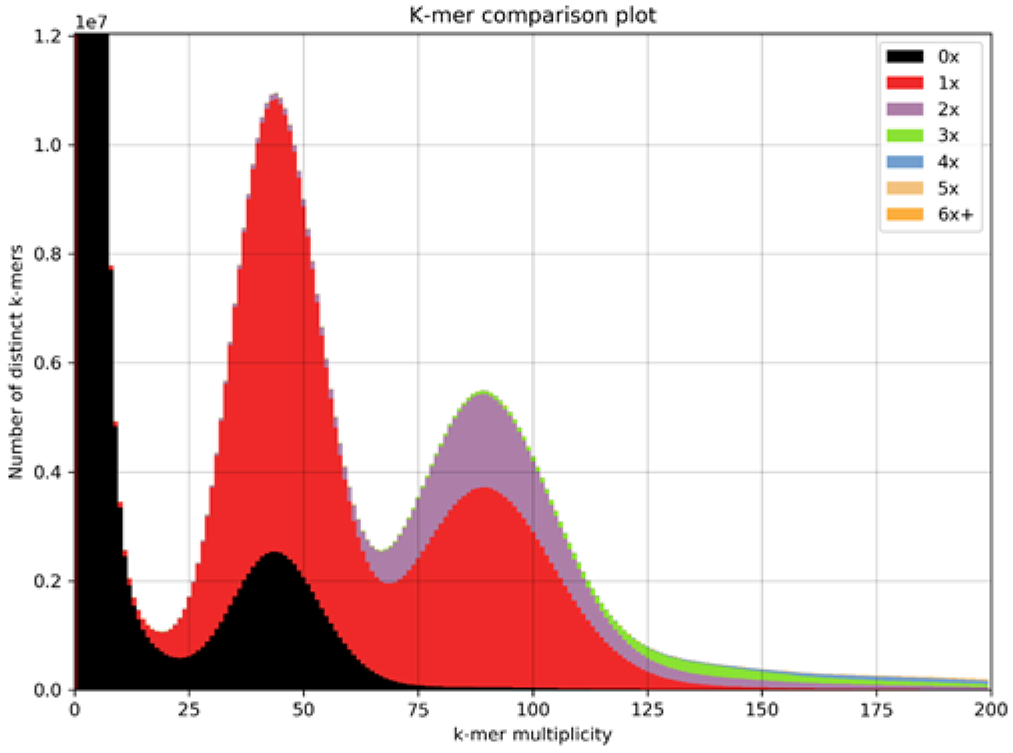


**B**

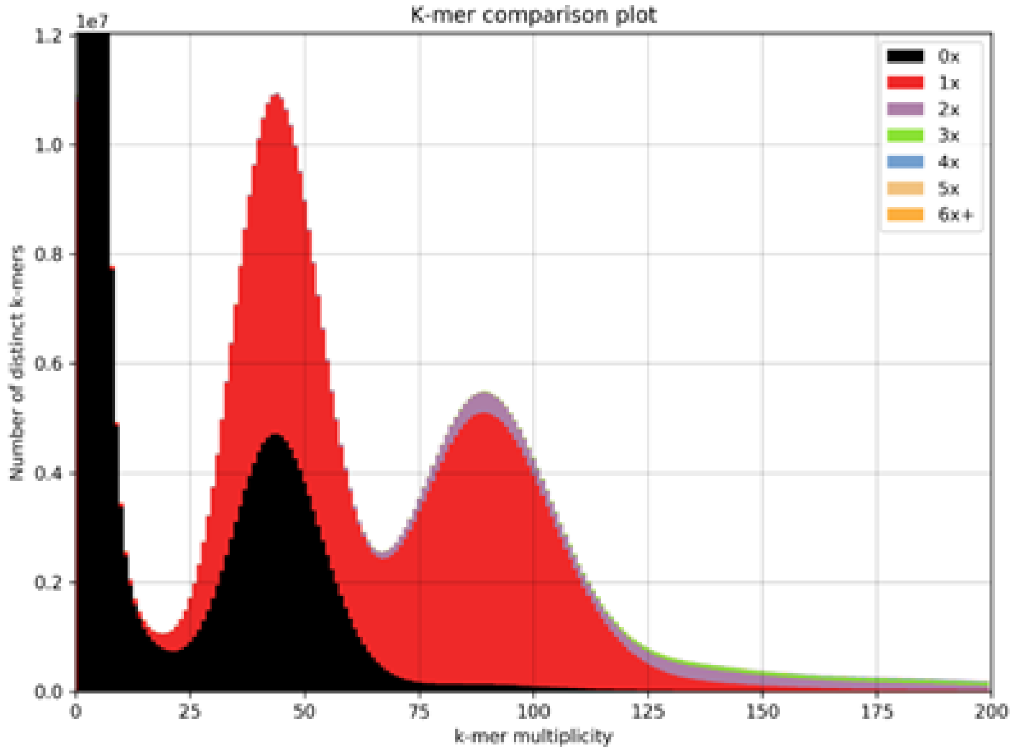

**Figure S3.** KAT plot of *Pocillopora meandrina* assemblies. (A) polished nanopore assembly. (B) haploid version of the genome.

**A**

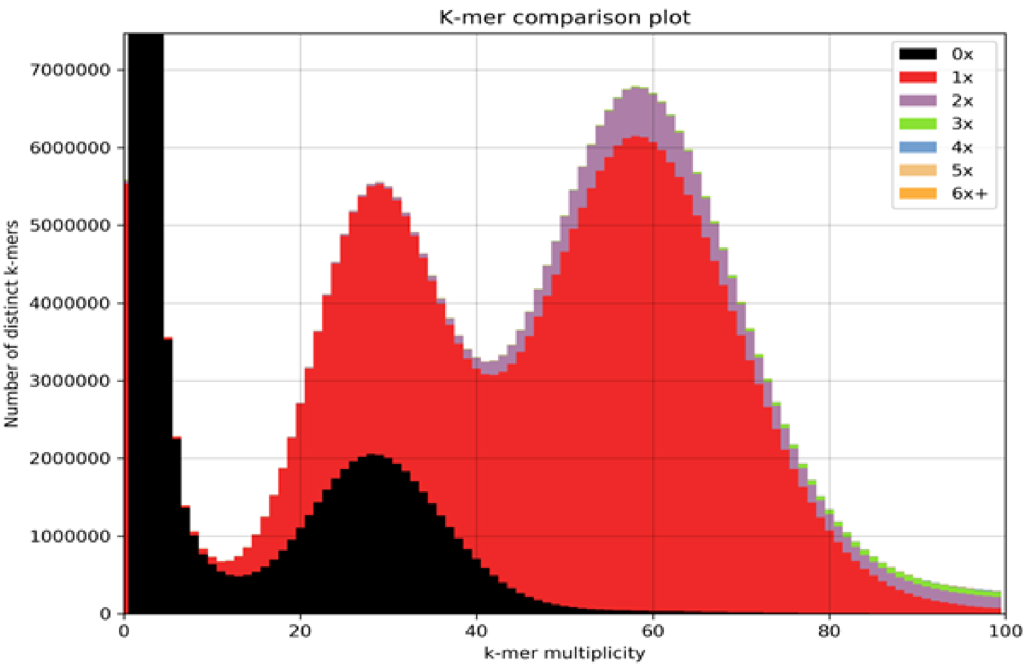


**B**

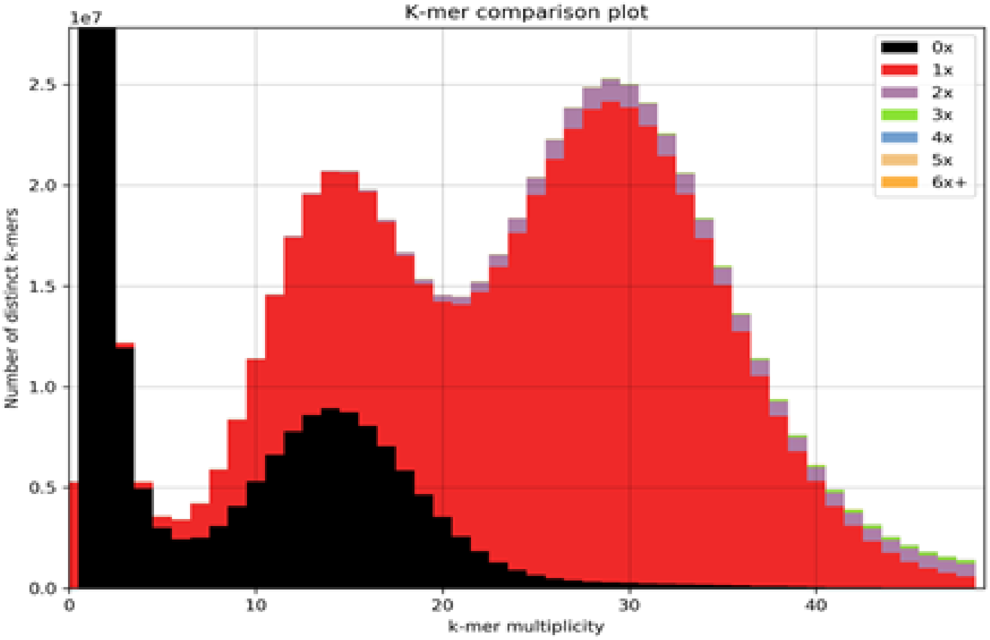

**Figure S4.** K-mer profiling for all assemblies of *Porites lobata* (a, b and c) and *Pocillopora meandrina* (d, e and f). From the Illumina sequences and assemblies, k-mer multiplicity was calculated. The x-axis is the k-mer multiplicity calculated from reads, and the numbers (0X, 1X, 2X, …) represent the k-mer multiplicity found in the corresponding assembly. K-mer multiplicity of 2 copies (blue curve) corresponds to duplicated contigs or overlaps in the assembly.

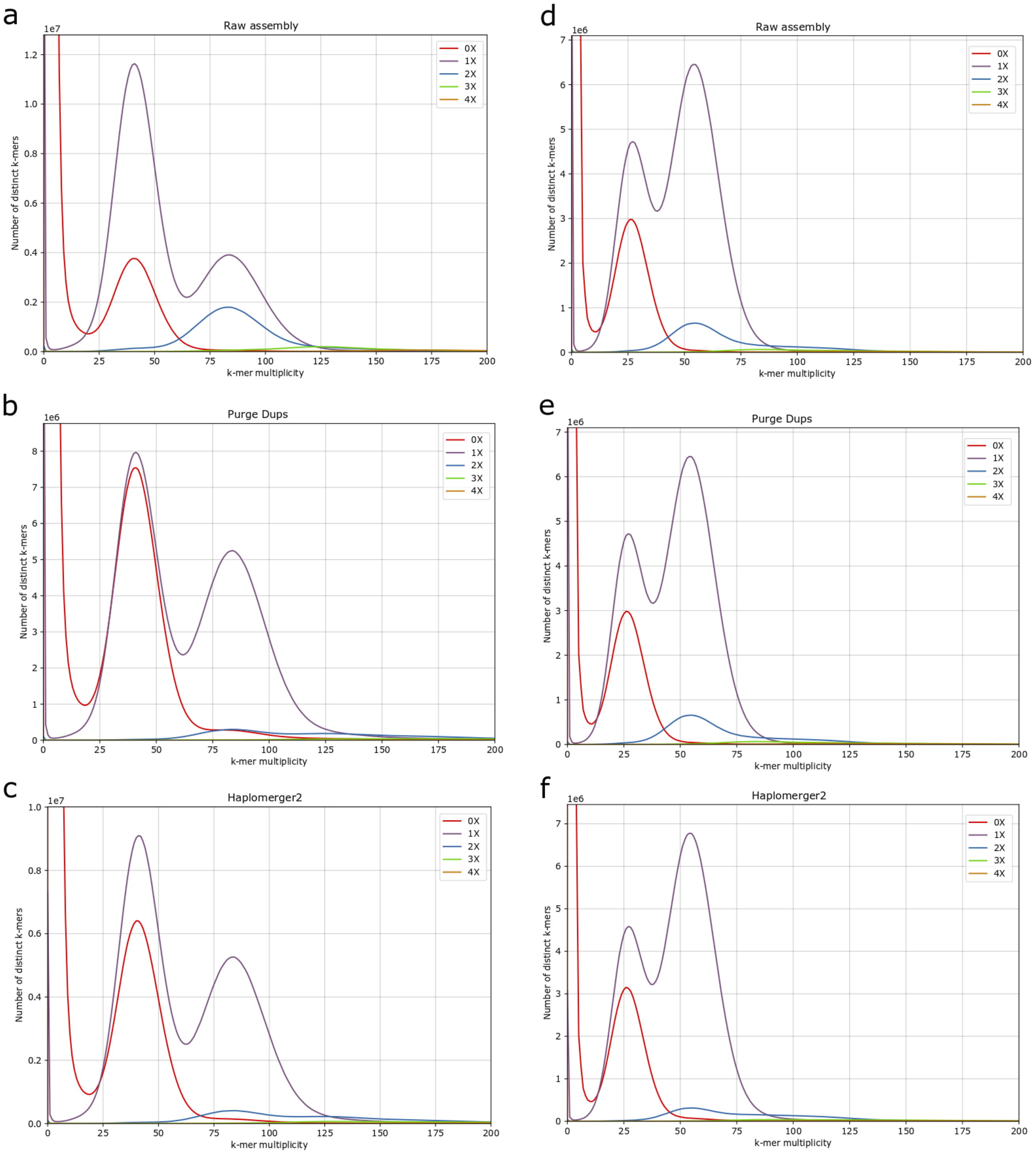

**Figure S5.** Distribution of the number of genes in tandemly repeated gene clusters for haplotype 1 and 2 assemblies generated by haplomerger2 for *Porites lobata* (A) and *Pocillopora meandrina* (B).

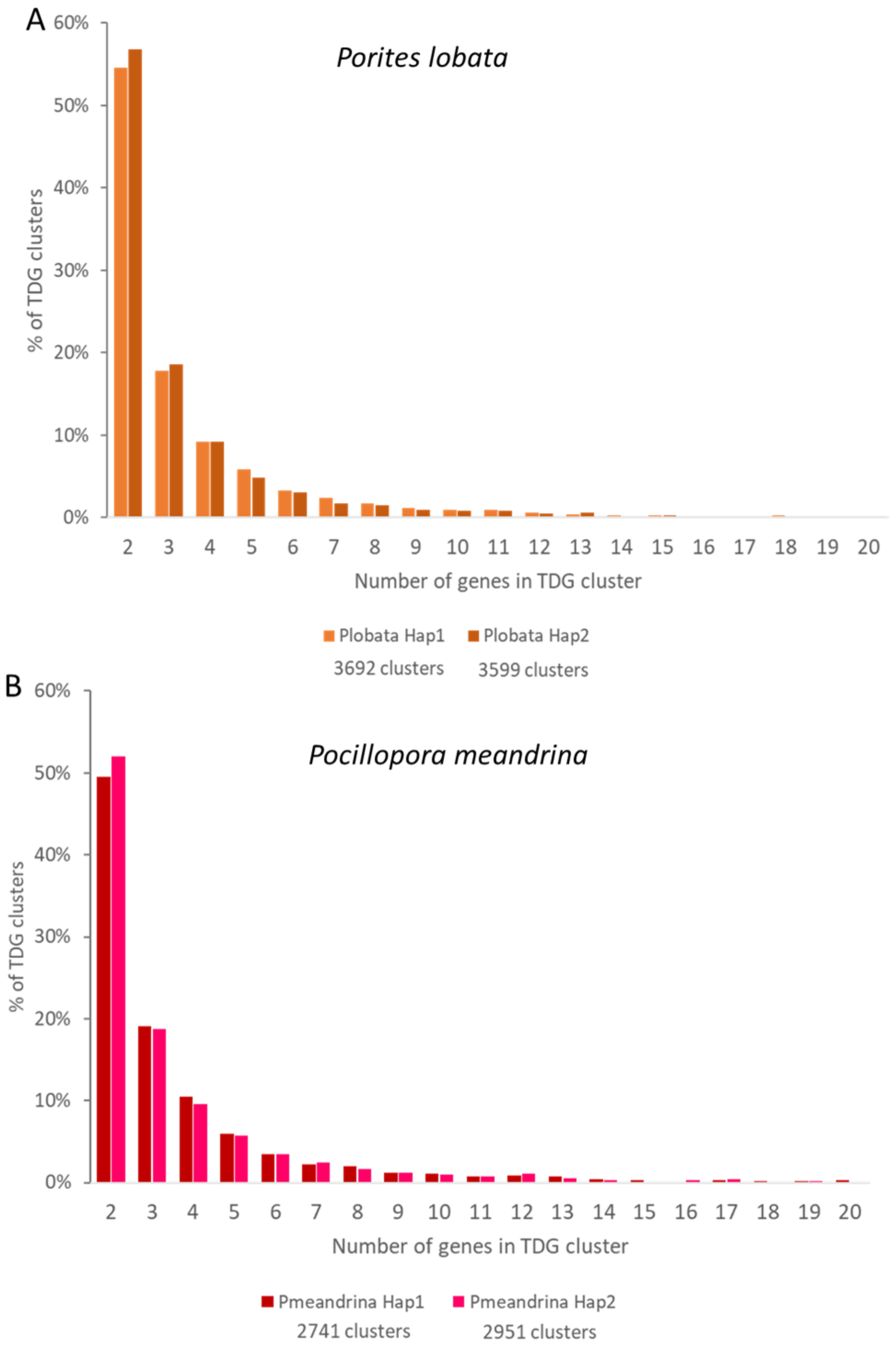

**Figure S6.** Validation of pairs of adjacent duplicated genes and clusters of TDG. Brown boxes represent three tandemly duplicated genes A, B and C, and blue rectangles represent ONT reads.

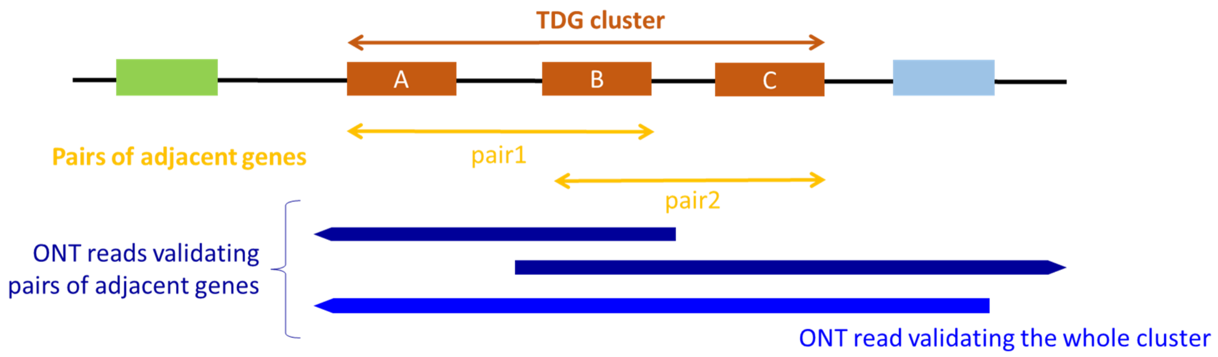

**Figure S7.** Validation of pairs of adjacent duplicated genes. Each dot represents a pair of two duplicated genes (**A**: *Porites lobata* N=8,220 and **B**: *Pocillopora meandrina* N=6,552) with x=0 for pairs that could not be validated by an ONT read and x=1 if at least one ONT read completely overlaps the two genes (Table S9). The y axis represents the distance from the start of the first gene to the end of the second gene. On the right panel, the read length distribution is given for comparison.

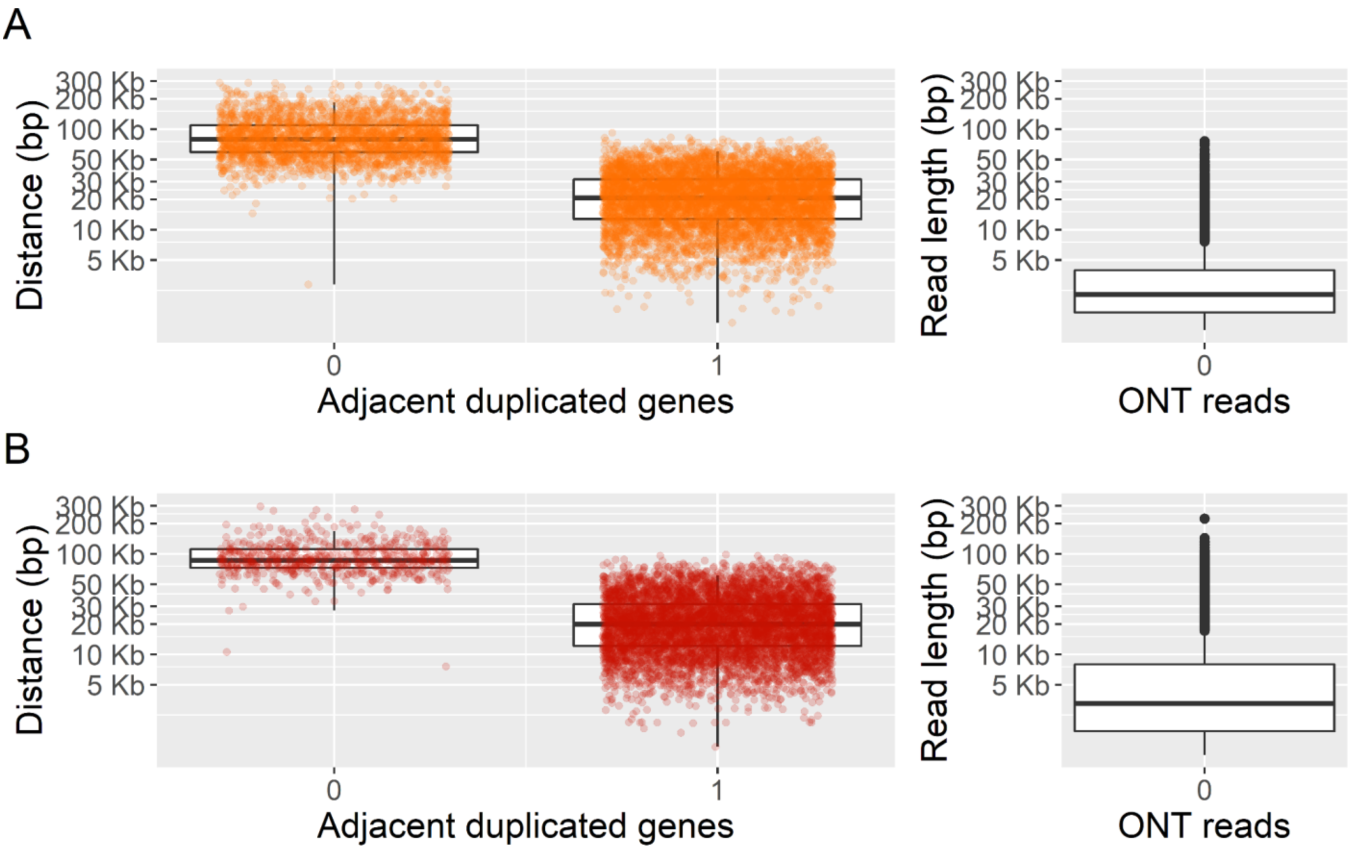

**Figure S8.** Validation of TDG clusters. Each dot represents a cluster of TDG (**A**: *Porites lobata* N=3,692 and **B**: *Pocillopora meandrina* N=2,741) with x=0 for clusters that could not be validated by an ONT read and x=1 if at least one ONT read completely overlaps all the genes of the cluster (Table S9). The y axis represents the distance from the start of the first gene to the end of the last gene. On the right panel, the read length distribution is given for comparison.

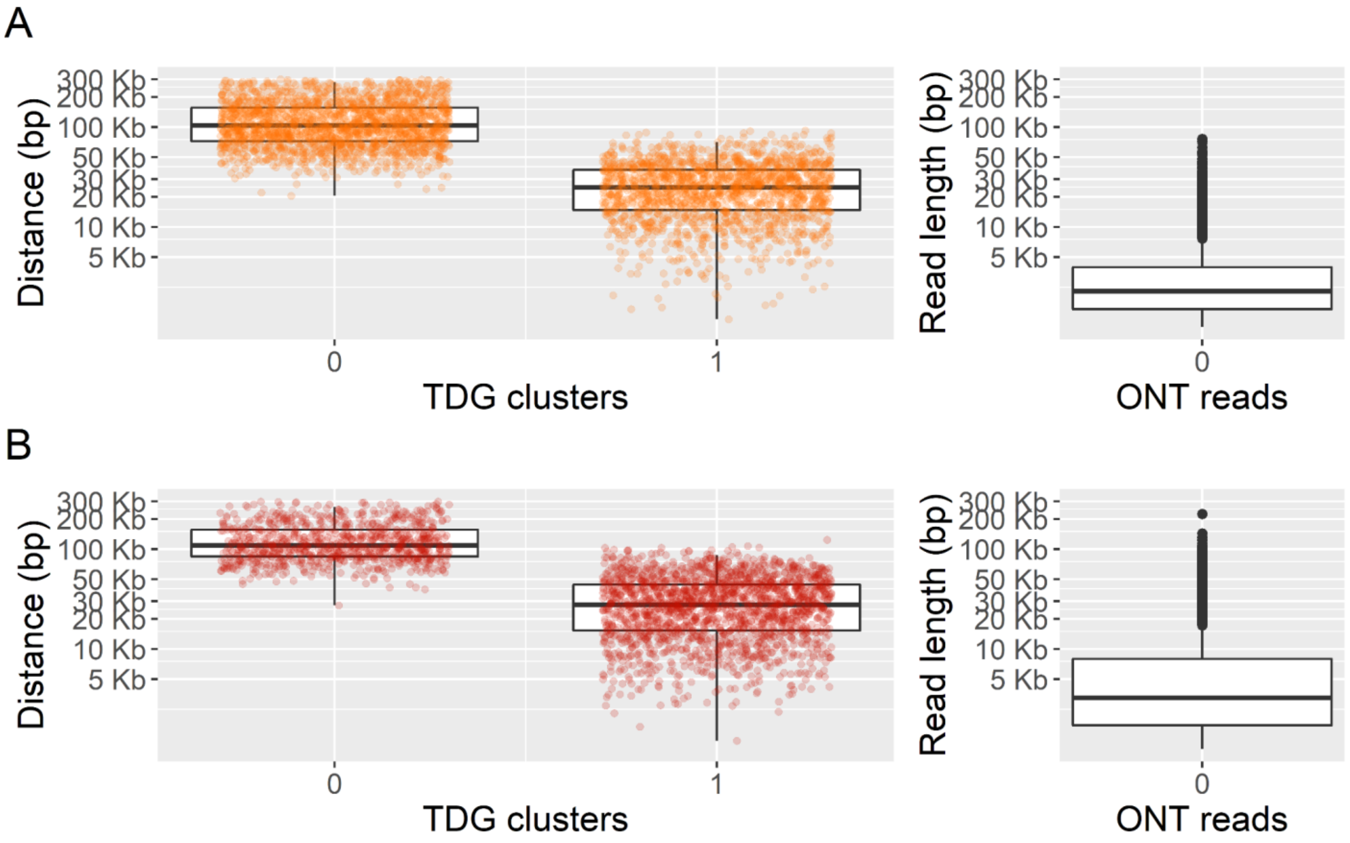

**Figure S9.** Comparison of haplotype assemblies and annotations for a TDG cluster (belonging to OG000308, amplified in corals, containing “GPCR” domain). **A**. Gbrowse view of Haplotype 1 (top) and 2 (bottom) assemblies. TDG cluster 3154 on haplotype 1 is validated by 10 nanopore reads spanning the whole cluster, and corresponds to TDG cluster 1055 on haplotype 2 (all gene pairs are BRH=best reciprocal hits). Ks between alleles are very low but some allelic differences are observed as seen in **B**. Zoom on an intronic region (gray triangle in A on Porites_lobata_contig_18:2,293,120..2,293,239) : allelic polymorphisms between haplotype 1 and 2 are validated by illumina and nanopore reads (one nanopore read is displayed for each haplotype). Reads are mapped on haplotype 1 and mismatches are displayed in yellow, deletions in red, insertions in purple.

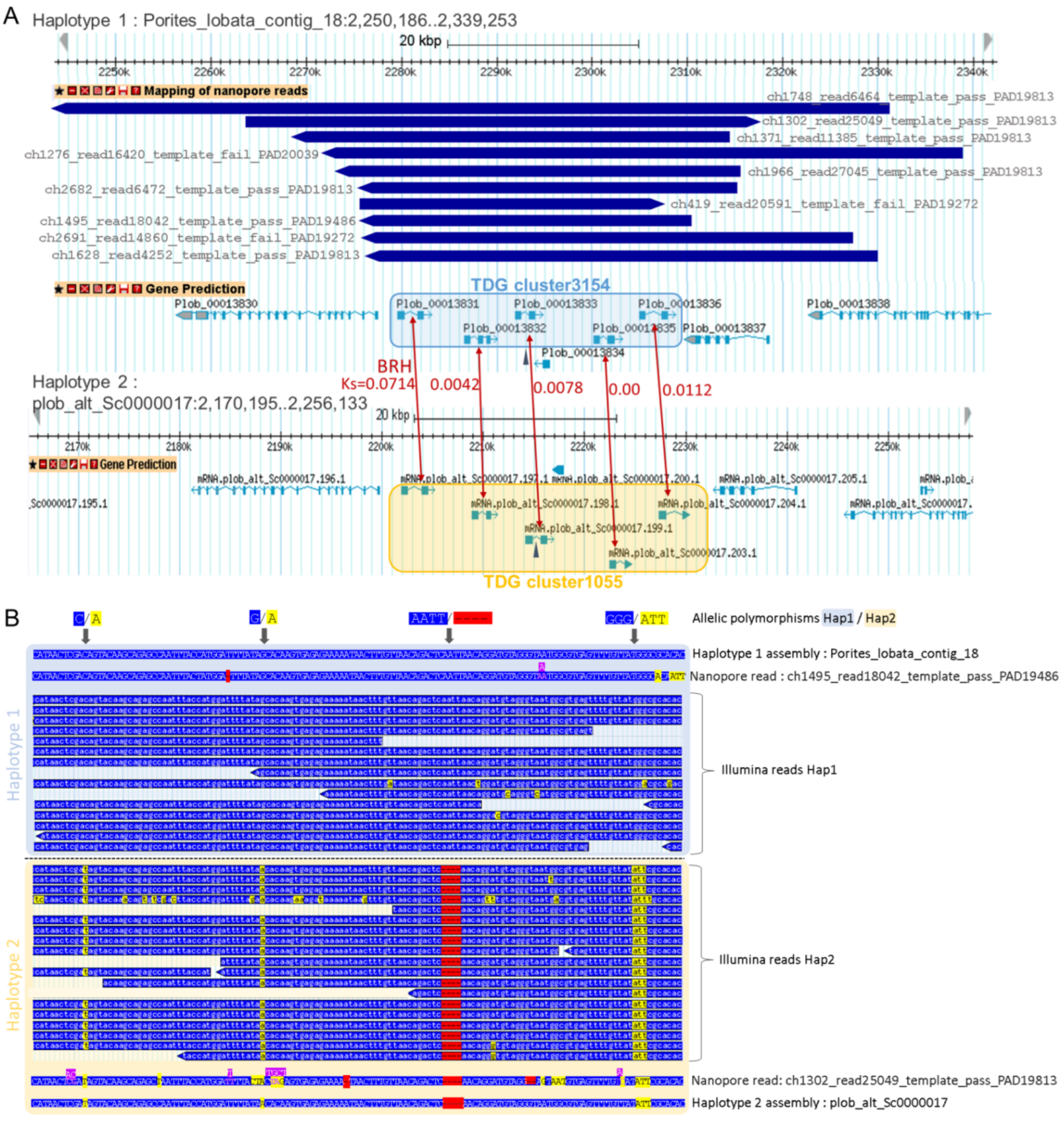

**Figure S10.** Two *P. lobata* genomic regions that illustrate proteins (A, green tracks) and transcripts (B, orange tracks) alignments before (middle track) or after (down track) the adaptation of gene prediction workflow for tandemly duplicated genes (“TDG patch”).

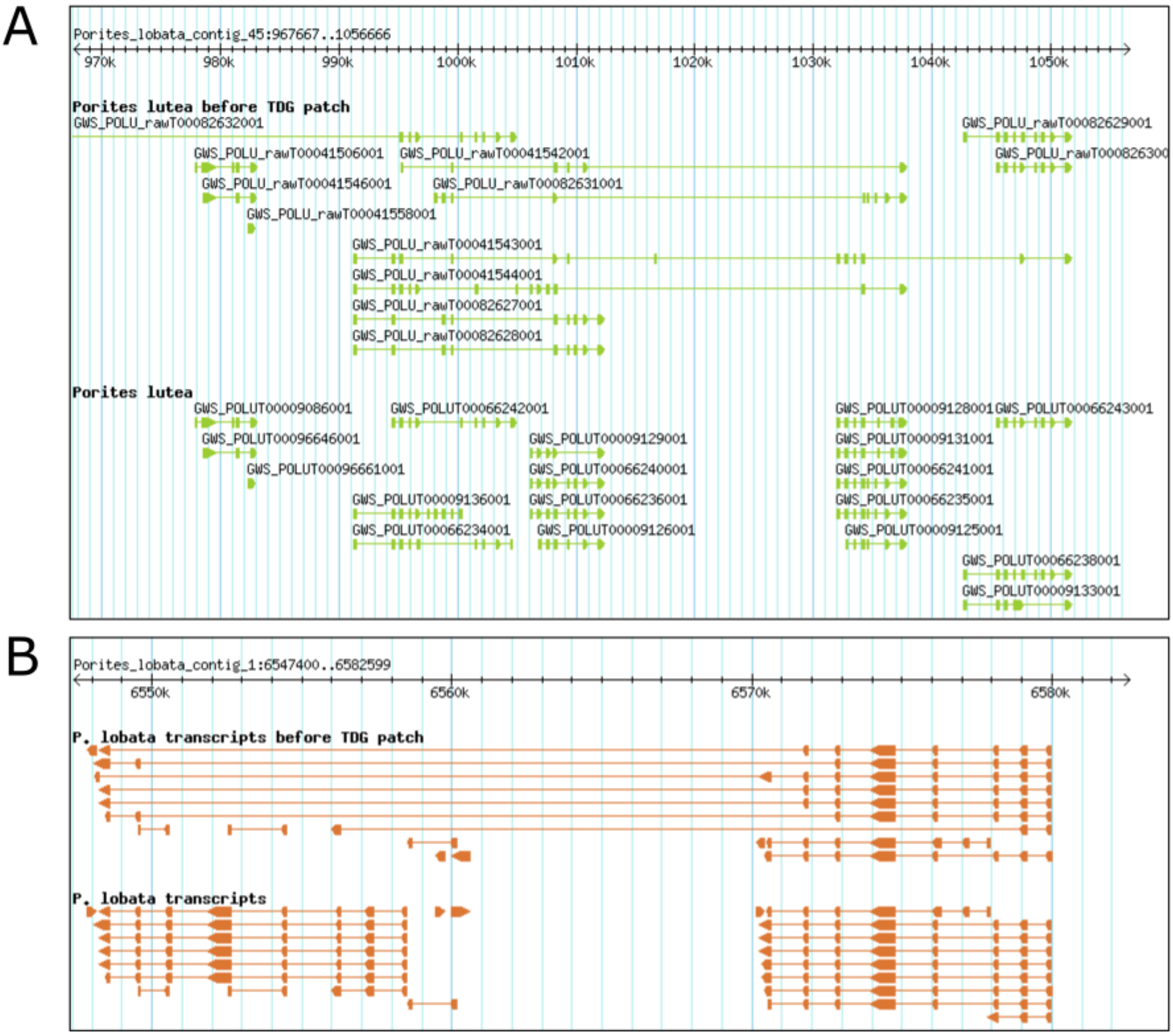

**Figure S11.** Consensus sequences of palindromic ITS satellites, found in *Porites lobata*, *Porites evermanni* and *Porites lutea* genomes. The telomeric palindrome is indicated by red arrows. These satellites are repeated in tandem in intergenic regions.

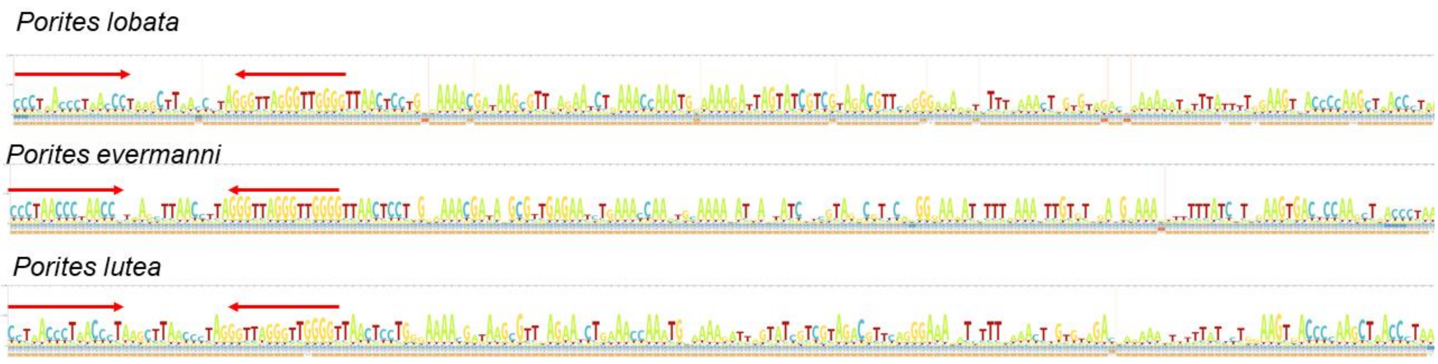

**Figure S12. Conservation of orthologous genes in *Porites lobata* and *Pocillopora meandrina* with other Cnidarian.** Distribution of identity percent between orthologs, grouped by clade.

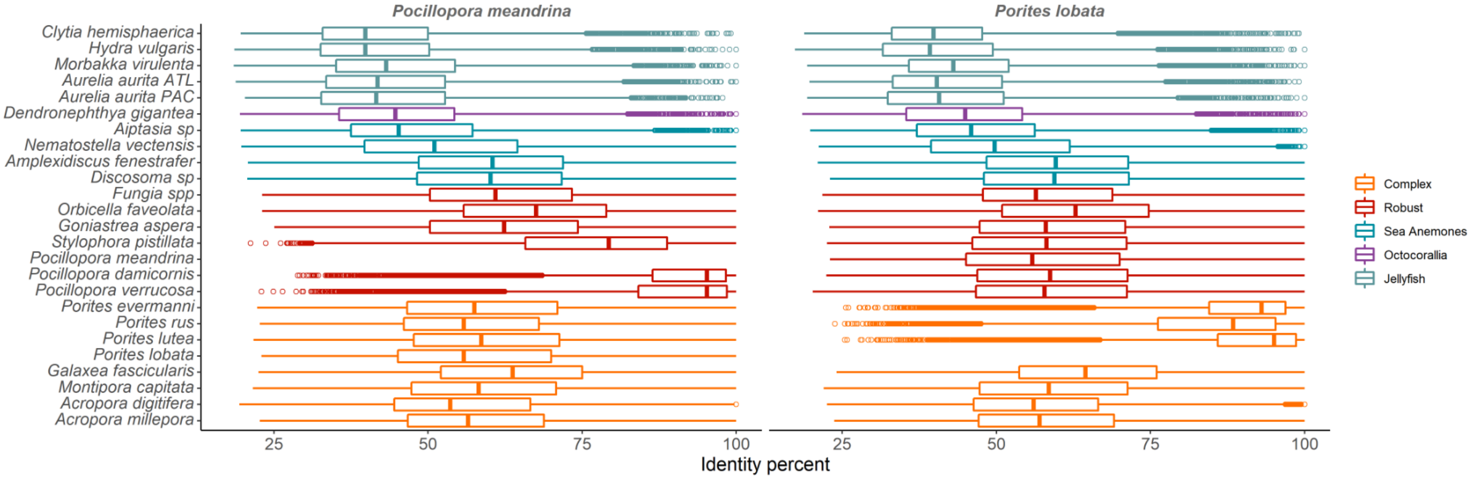


**Figure S13. Number of TDG genes vs total number of genes on contigs.** Total number of genes (x axis) vs number of TDG (with Ks<0.5) (y axis) for contigs with 10 genes or more in *Porites lobata (A)* and *Pocillopora meandrina* (B). Each dot represents a contig.

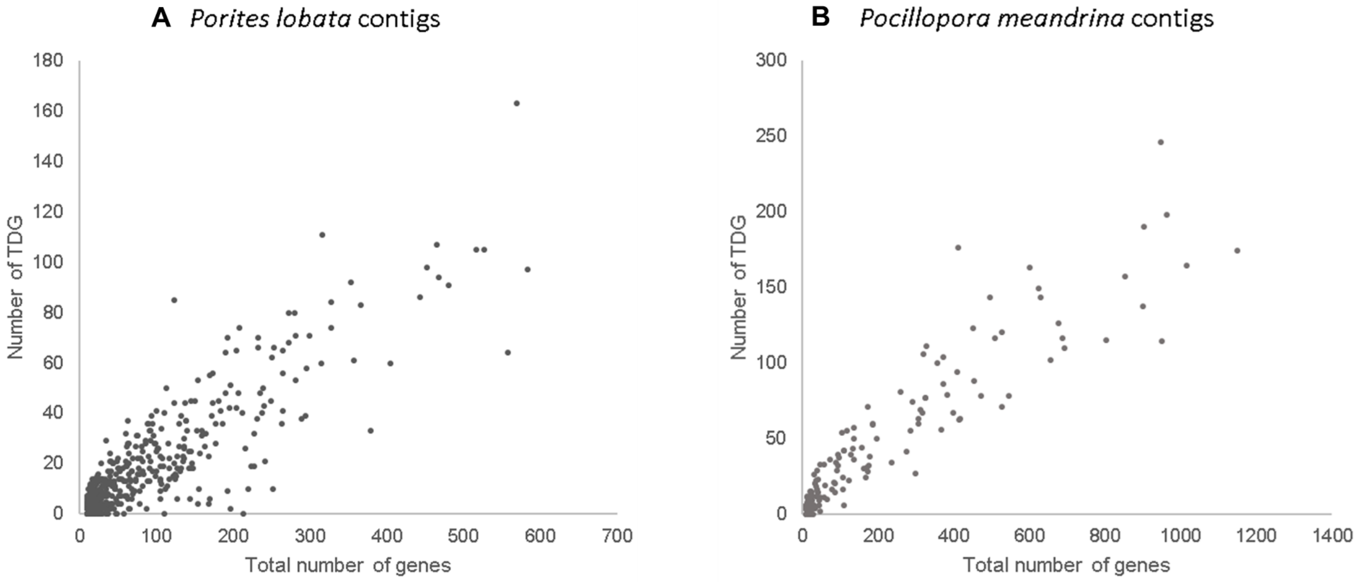

**Figure S14**. **Expanded PFAM domains in corals vs non-corals hexacorallia.**

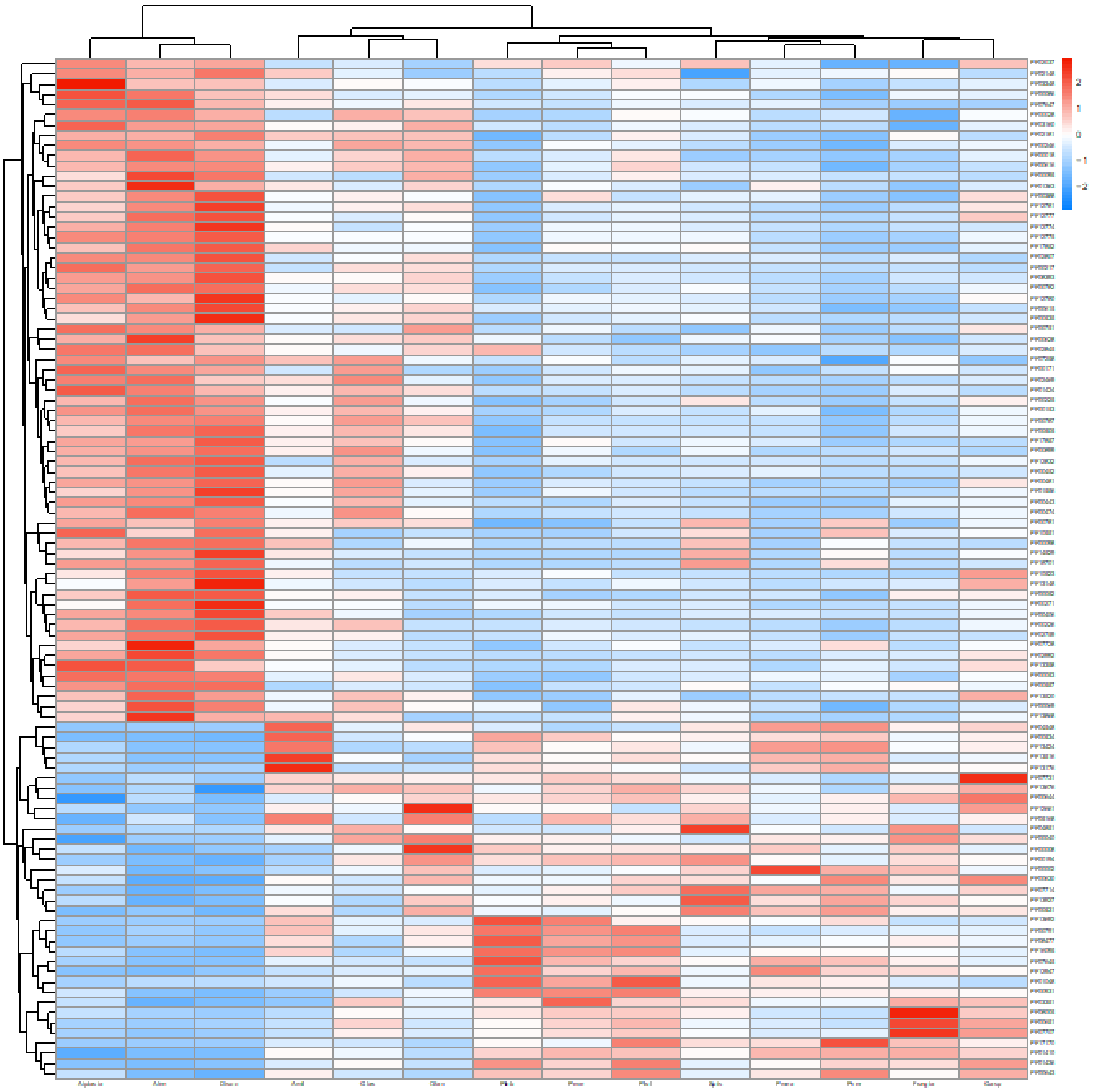

**Figure S15. OGs amplified in complex vs robust.** Heatmap representing the number of genes belonging to each OG.Complex and robust species are respectively grouped in the orange and dark red lines in the phylogenetic tree.

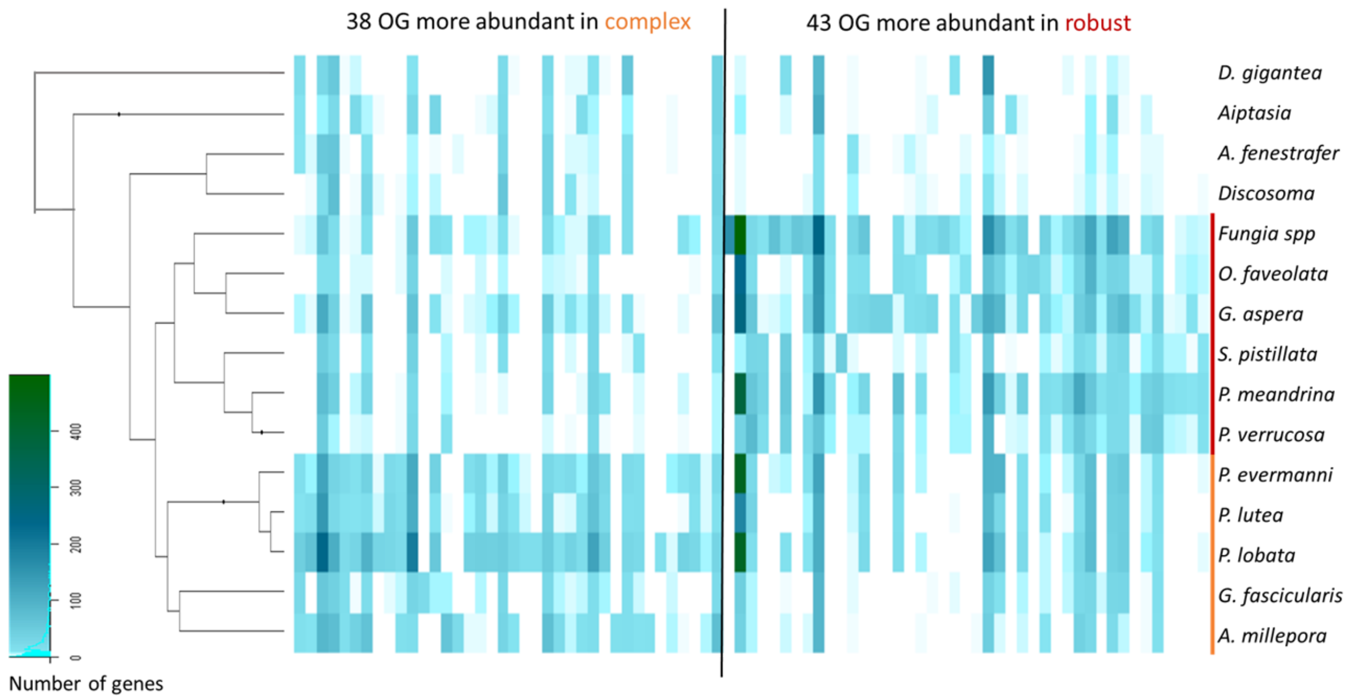

**Figure S16. Expanded PFAM domains in complex vs robust corals.**

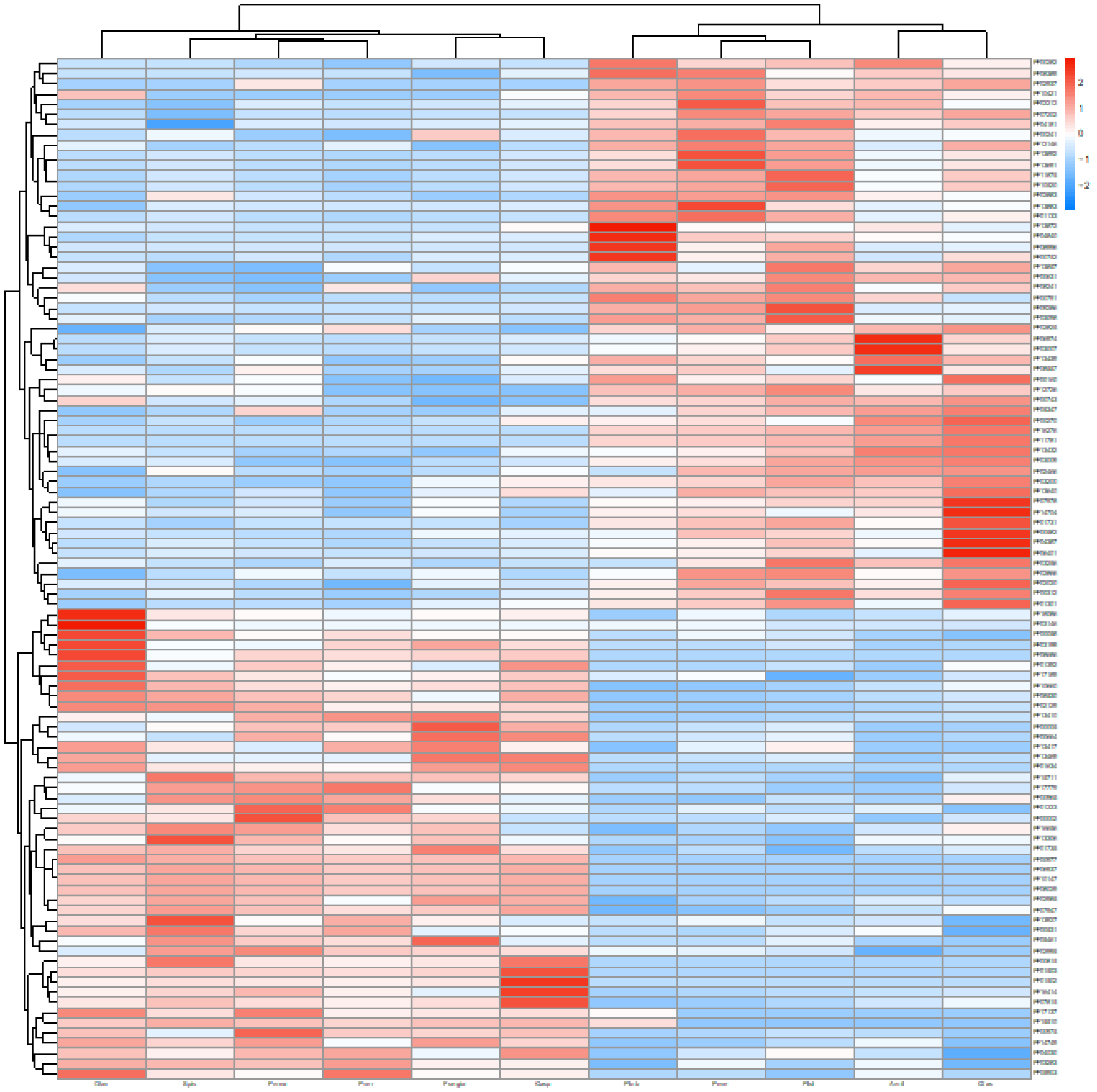

**Figure S17. OGs amplified in massive vs branched coral species.** Heatmap representing the number of genes belonging to each OG. Green and purple lines show species belonging to massive and branched colonies (respectively). Species names are followed by C when belonging to the complex clade and R when belonging to the robust clade.

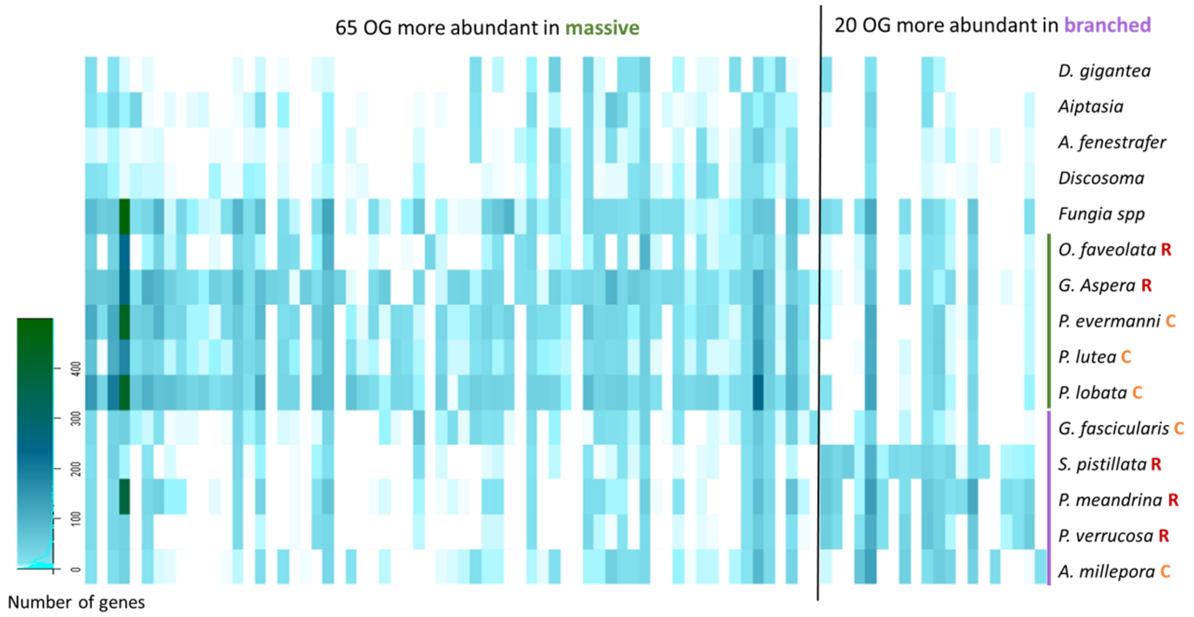

**Figure S18.** Heatmap representing the P-values obtained with CAFE for gene number amplification (in red) and reduction (in blue) events for each node on the phylogenetic tree.

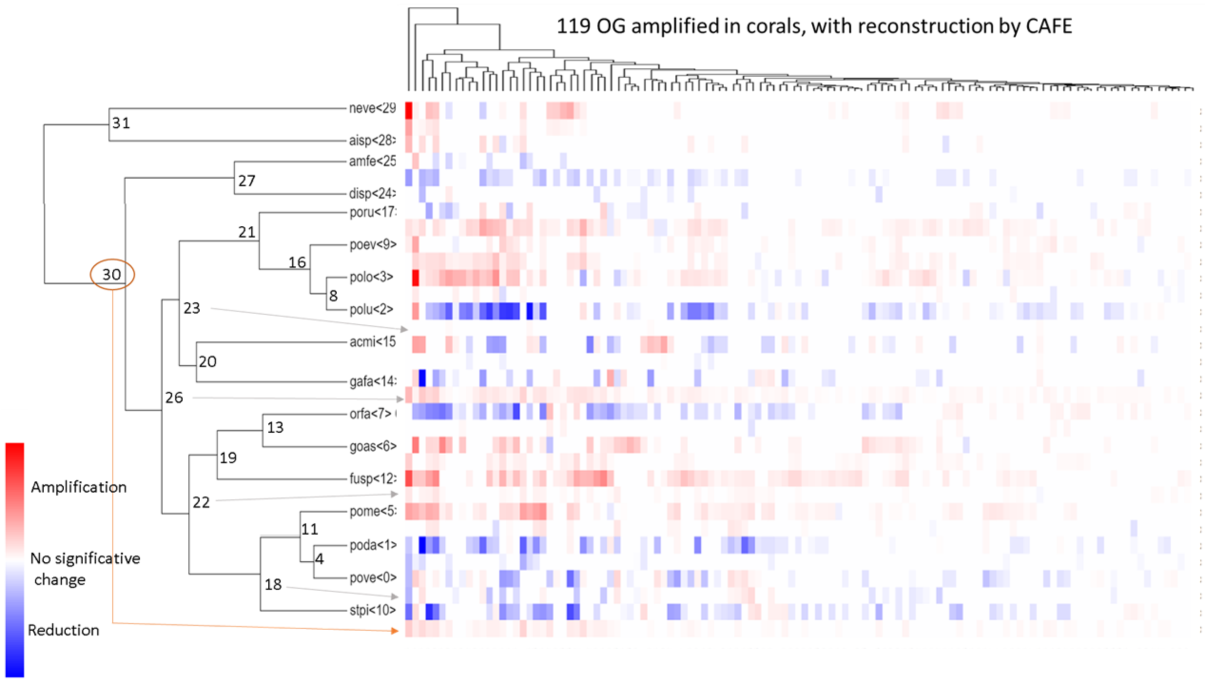

**Figure S19. Gene family amplifications/reductions identified with CAFE.** Results of CAFE reconstructions on gene families (orthogroups) present at the root of Hexacorallia (corals + sea anemones), in 17 species. Species names are the following: neve= Nematostella vectensis, aisp= *Aispasia* sp, amfe= *Amplexidiscus fenestrafer*, disp= *Discosoma* sp, poru= *Porites rus*, poev= *Porites evermanni*, polo= *Porites lobata*, polu= *Porites lutea*, acmi= *Acropora millepora*, gafa= *Galaxea fascicularis*, orfa= *Orbicella faveolata*, goas= *Goniastrea aspera*, fusp= *Fungia* sp, pome= *Pocillopora meandrina*, poda= *Pocillopora damicornis*, pove= *Pocillopora verrucosa*, stpi= *Stylophora pistillata*. **A.** phylogenetic tree constructed by orthofinder, with branches colored according to the number of significant gene family amplifications detected by CAFE, from white (0) to red (383, in the final branch leading to *Porites lobata*). **B.** Barplot representing the number of orthogroups (OG) with significant gene number variation for each species (all events that occured on the branches leading to the species are taken into account). The number of amplified gene families (OG) are in red and the number of contracted gene families (OG) are in blue.

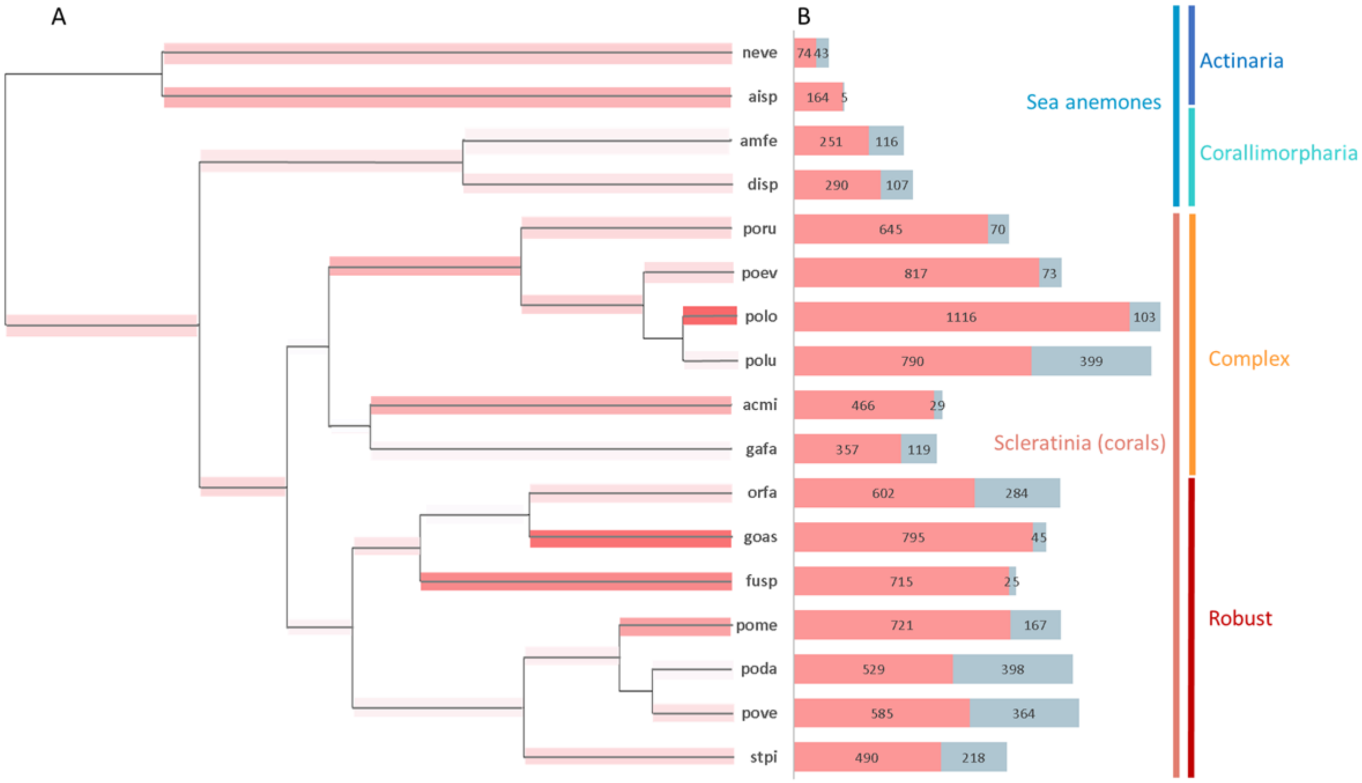

**Figure S20. Distribution of Ks between TDG gene pairs in four classes of intergenic distances.**

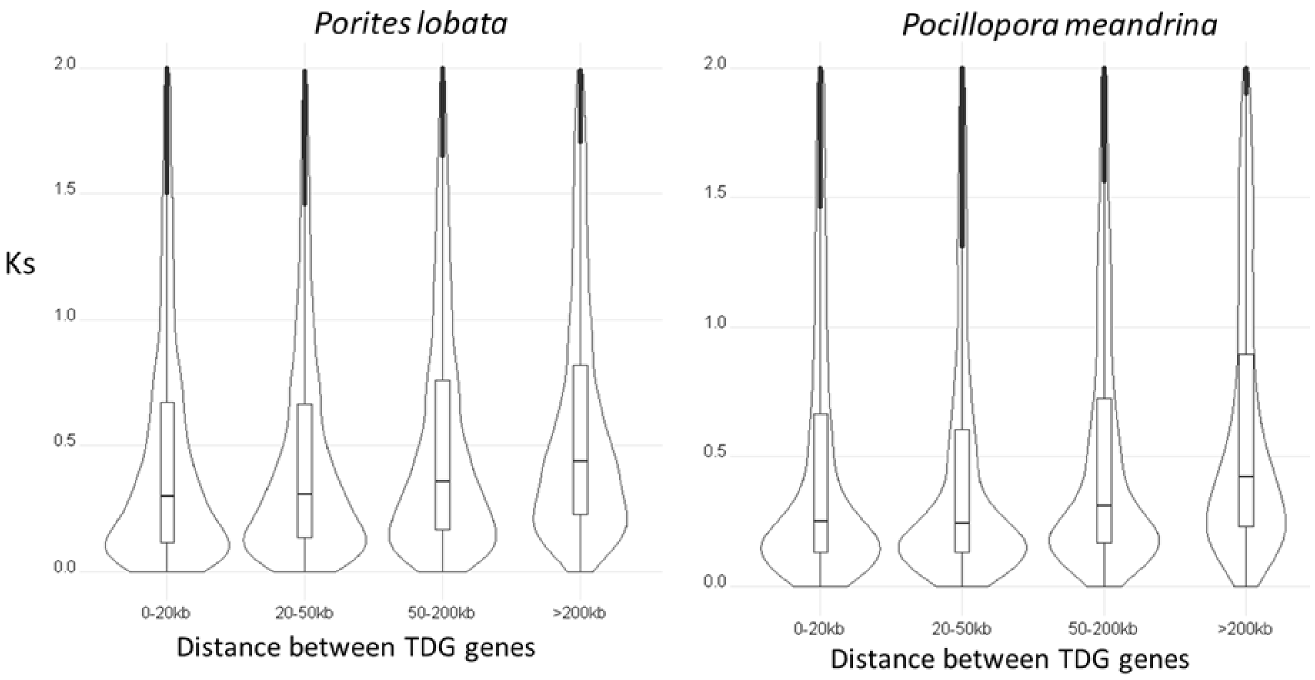


**Figure S21. View of the *Pocillopora meandrina* genome browser, with 2 adjacent genes duplicated in tandem.**

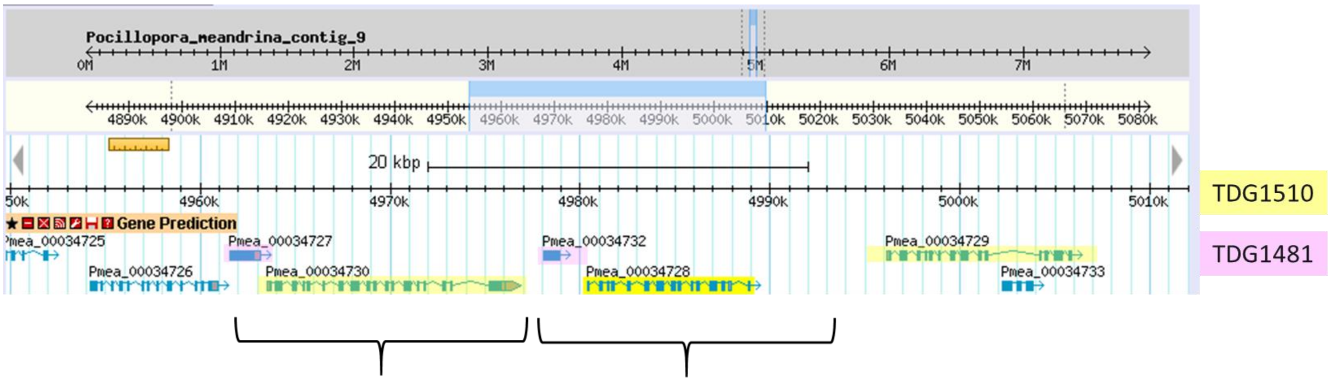

**Figure S22. Distribution of Ks between TDG pairs in 3 structural conservation classes** ("conserved" exons or introns is relative to the structure: it means that they have exactly the same size)**.** Numbers of pairs in each category are mentioned with "N=".

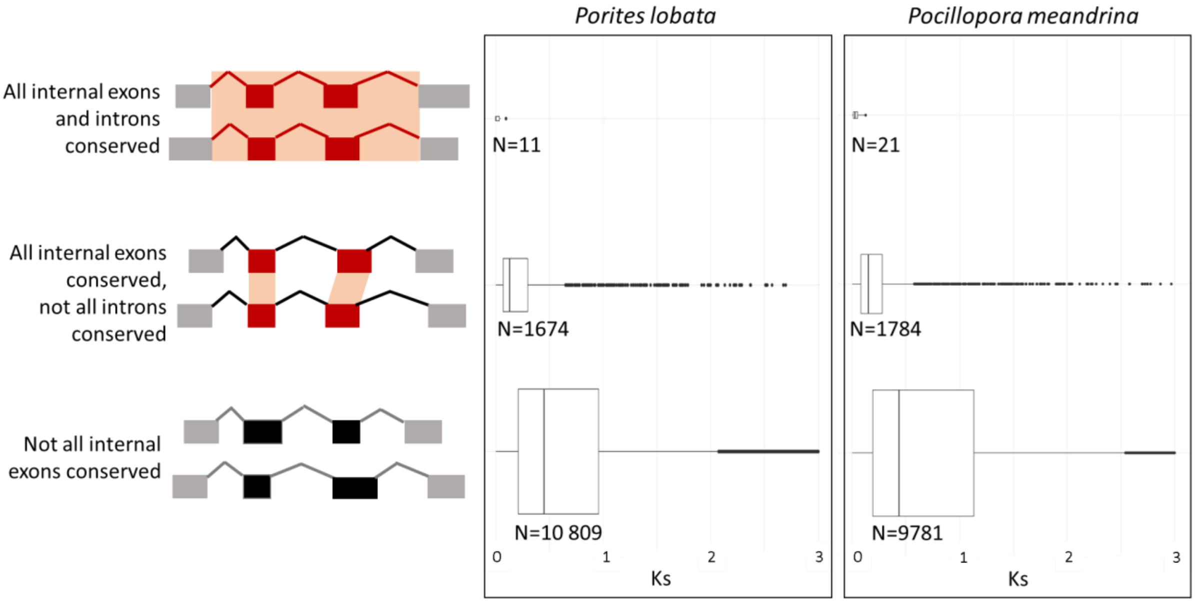

**Figure S23.** Browser view of a cluster of 3 tandemly duplicated genes belonging to OG0000023 (lectin-like domain). **B.** Structural comparison between the three annotated genes. **C.** Multiple alignment of 2 exons and one intron showing higher homology in exons.

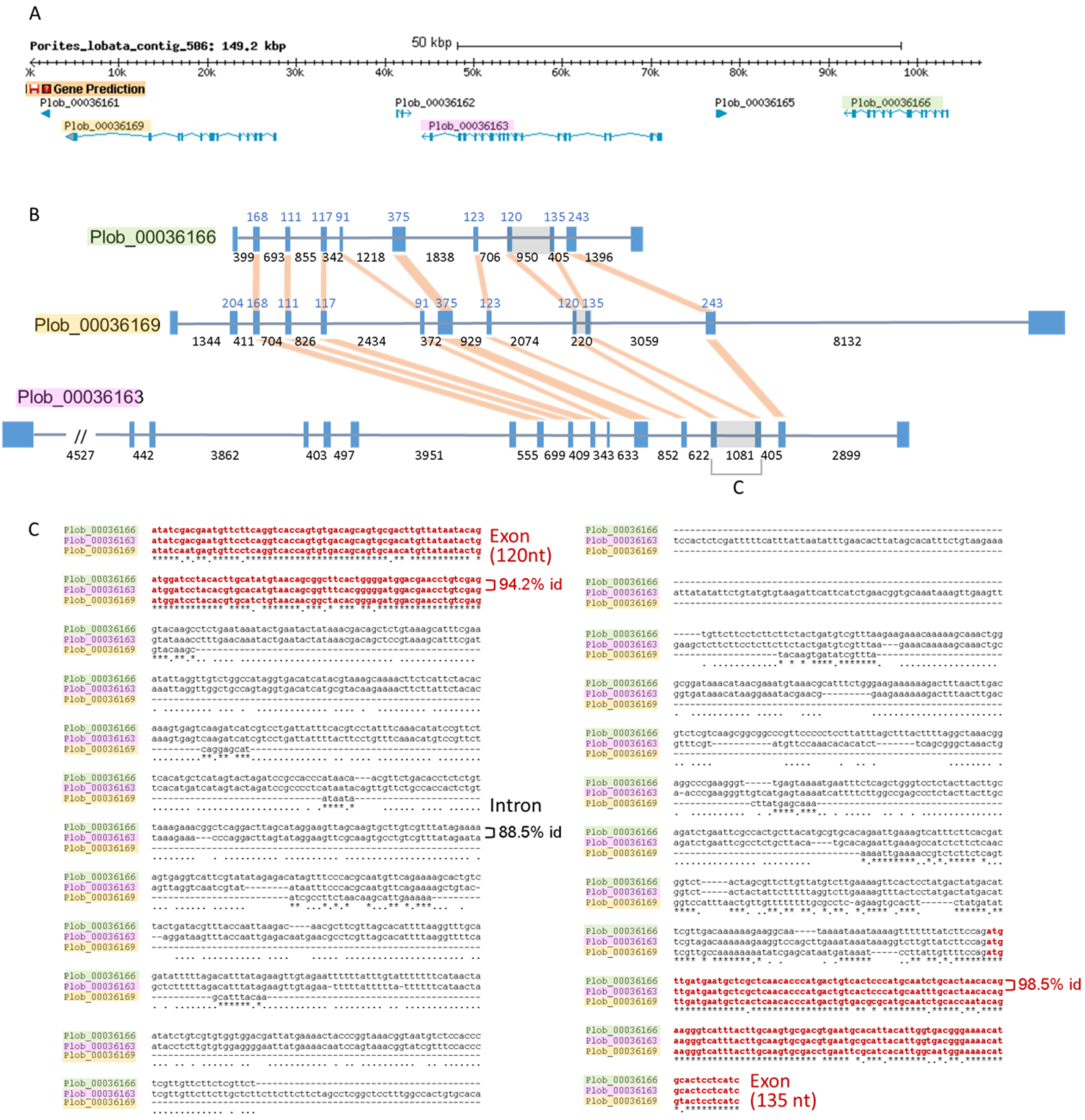

**Figure S24. Expression quantification of 200 random OGs across the 103 meta-transcriptome samples.** The heatmap shows the z-score of average TPM per OG (rows) according to the samples (columns). The z-score is computed by OG (row) using the scale function of the heatmap.2 from the gplots R package.

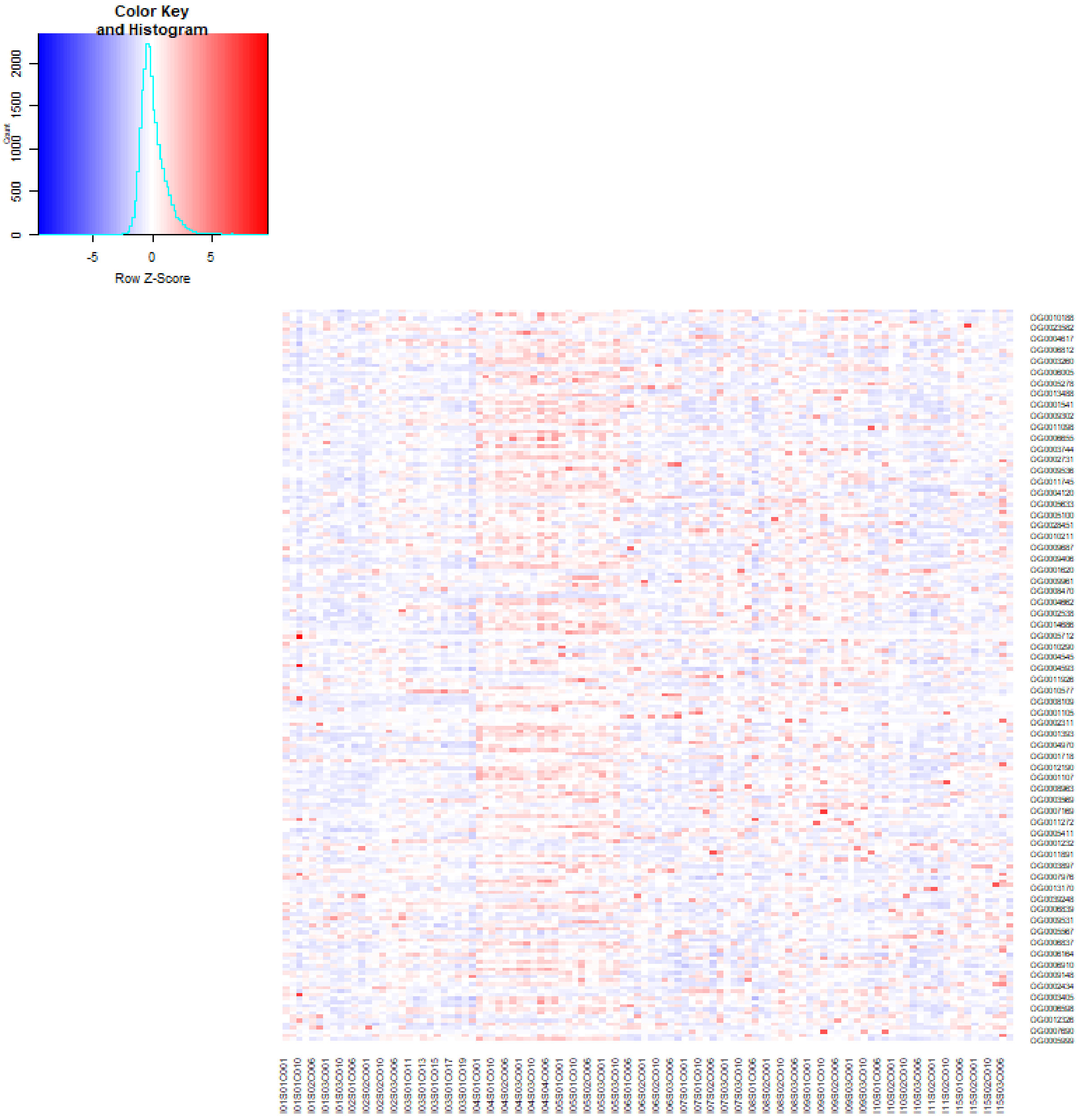

**Figure S25. Number of genes in OG identified as TLR (A), NLR (B), lectins (C).** Heatmaps (left panels) represent the number of genes in OGs for 14 cnidarian species. Histograms (right panels) [illegible]G.

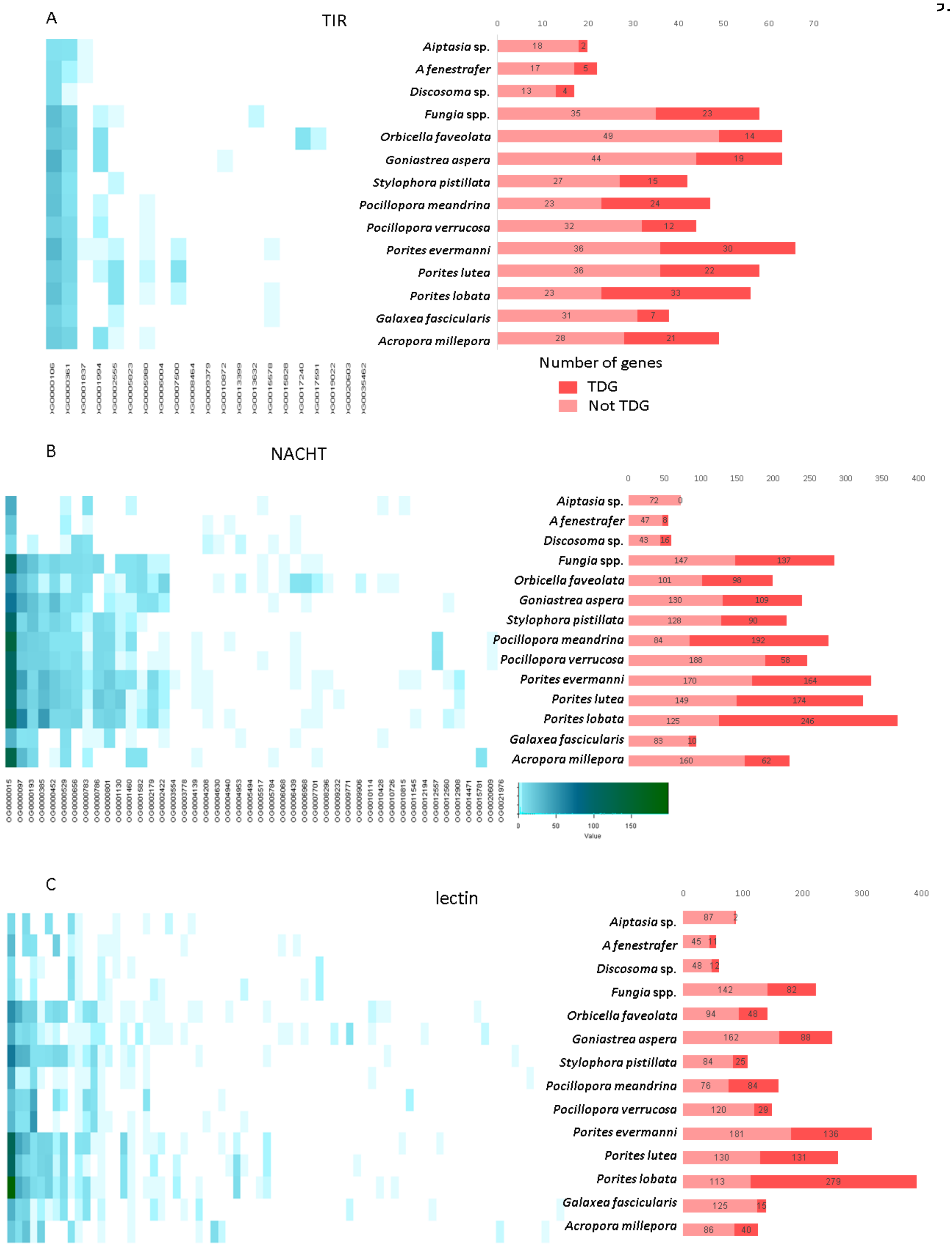

**Figure S26. Percent of NACHT/NB-ARC containing proteins that are likely truncated** at the N-terminal, C-terminal or both extremities (less than 250 aa upstream/downstream of the NACHT/NB-ARC domain), in 5 coral species. Total numbers of proteins are displayed with “N=”.

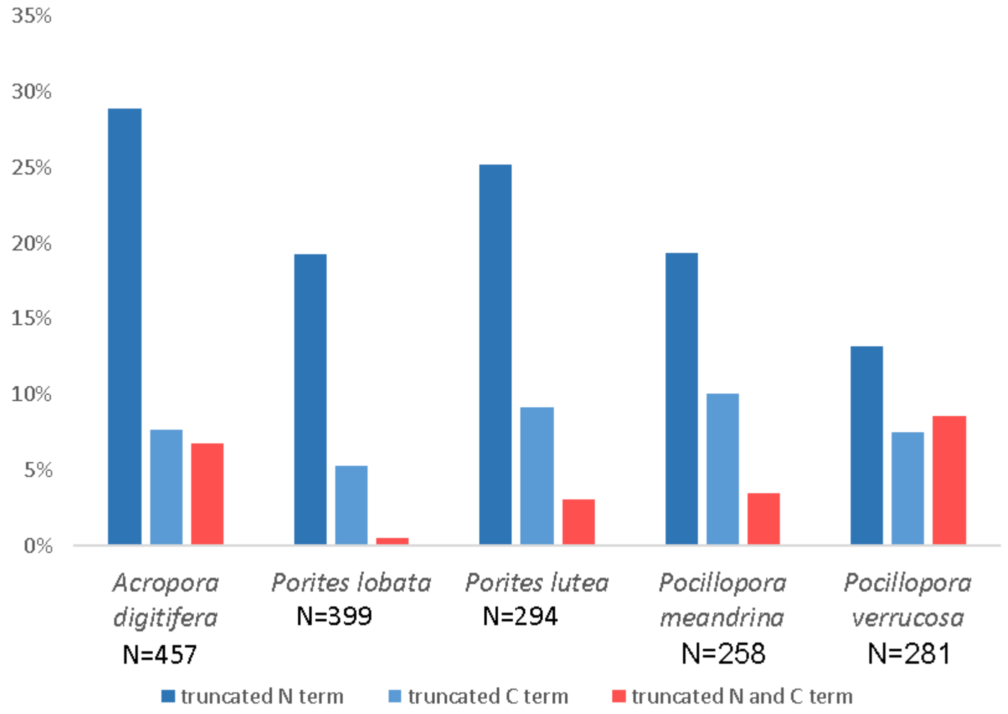

**Figure S27.** Observed N-terminal and C-terminal domains for NACHT/NB-ARC containing proteins in five coral species.

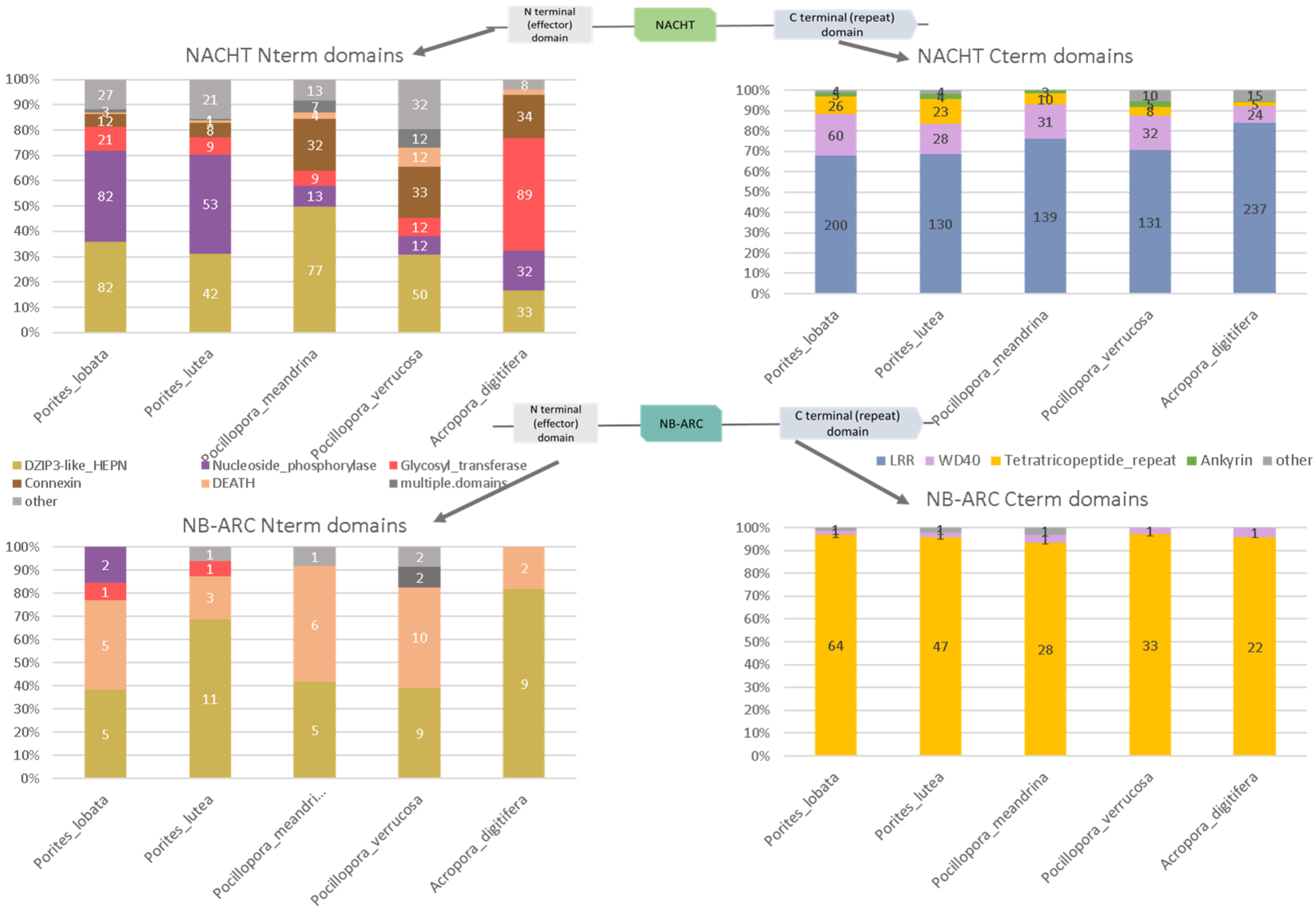

**Figure S28. A.** Graphic representation of the number of calcification-related proteins in non-scleractinian and scleractinian species **B.** *p*-values of calcification related proteins in non-scleractinian, scleractinian - robust and - complex coral species.

A.

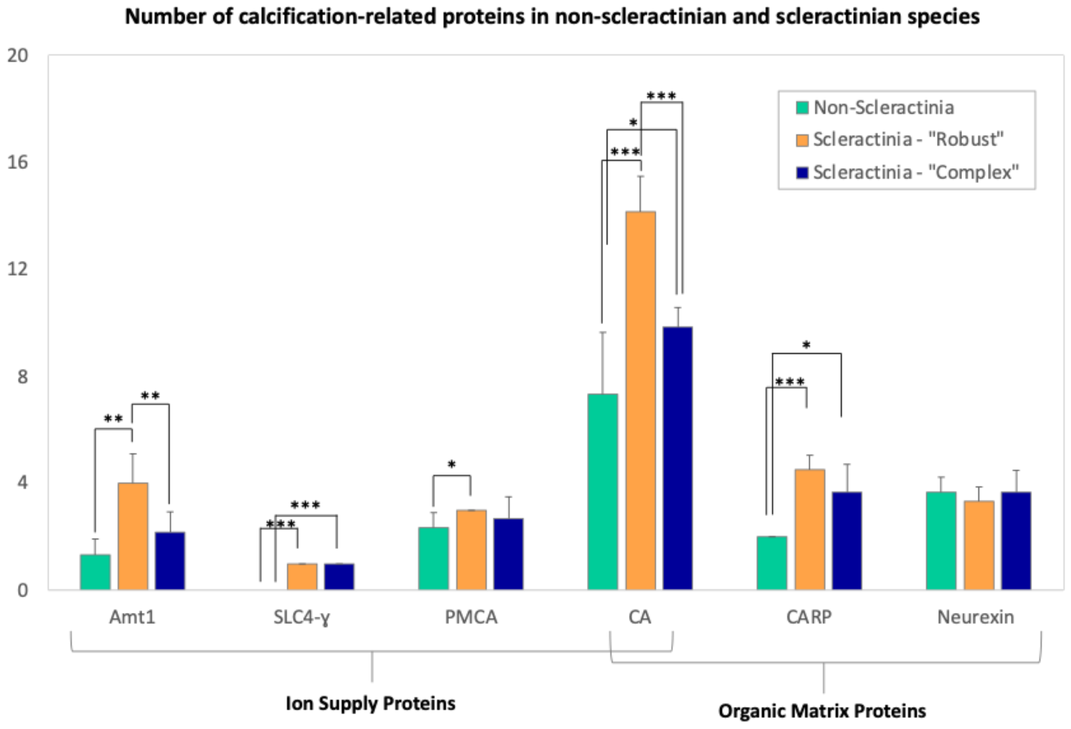


B.

| *p-value* | Non-Scleractinia vs Scleractinia - Robust | Non-Scleractinia vs Scleractinia - Complex | Scleractinia - Robust vs Scleractinia - Complex |
|---|---|---|---|
| **Amt1** | 0.00618 ** | 0.14 | 0.00702 ** |
| **SLC4-γ** | <2e-16 *** | <2e-16 *** | 0.341 |
| **PMCA** | 0.0185 * | 0.553 | 0.341 |
| **CA** | 0.00067 *** | 0.0383 * | 0.0000395 *** |
| **CARP** | 0.000122 *** | 0.0306 * | 0.111 |
| **Neurexin** | 0.407 | 1 | 0.418 |

**Figure S29. Maximum likelihood phylogenetic tree (Phyml, LG + I + G) of cnidarian ammonium transporters Amt1.** Cnidaria species include Corallimorpharia (*Discosoma sp.* - Dsp and *Amplexidiscus fenestrafer* - Afe), Actiniaria (*Exaiptasia diaphana* - Edi), Scleractinia - “Robust” (*Stylophora pistillata* - Spi, *Pocillopora verrucosa* - Pve, *Pocillopora meandrina* - Pme, *Orbicella faveolata* - Ofa, *Goniastrea aspera* - Gas, *Fungia sp.* - Fun) and Scleractinia - “Complex” (*Acropora digitifera* - Adi, *Acropora millepora* - Ami, *Galaxea fascicularis* - Gfa, *Montipora capitata* - Mca, *Porites lobata* - Plo, *Porites lutea* - Plu).

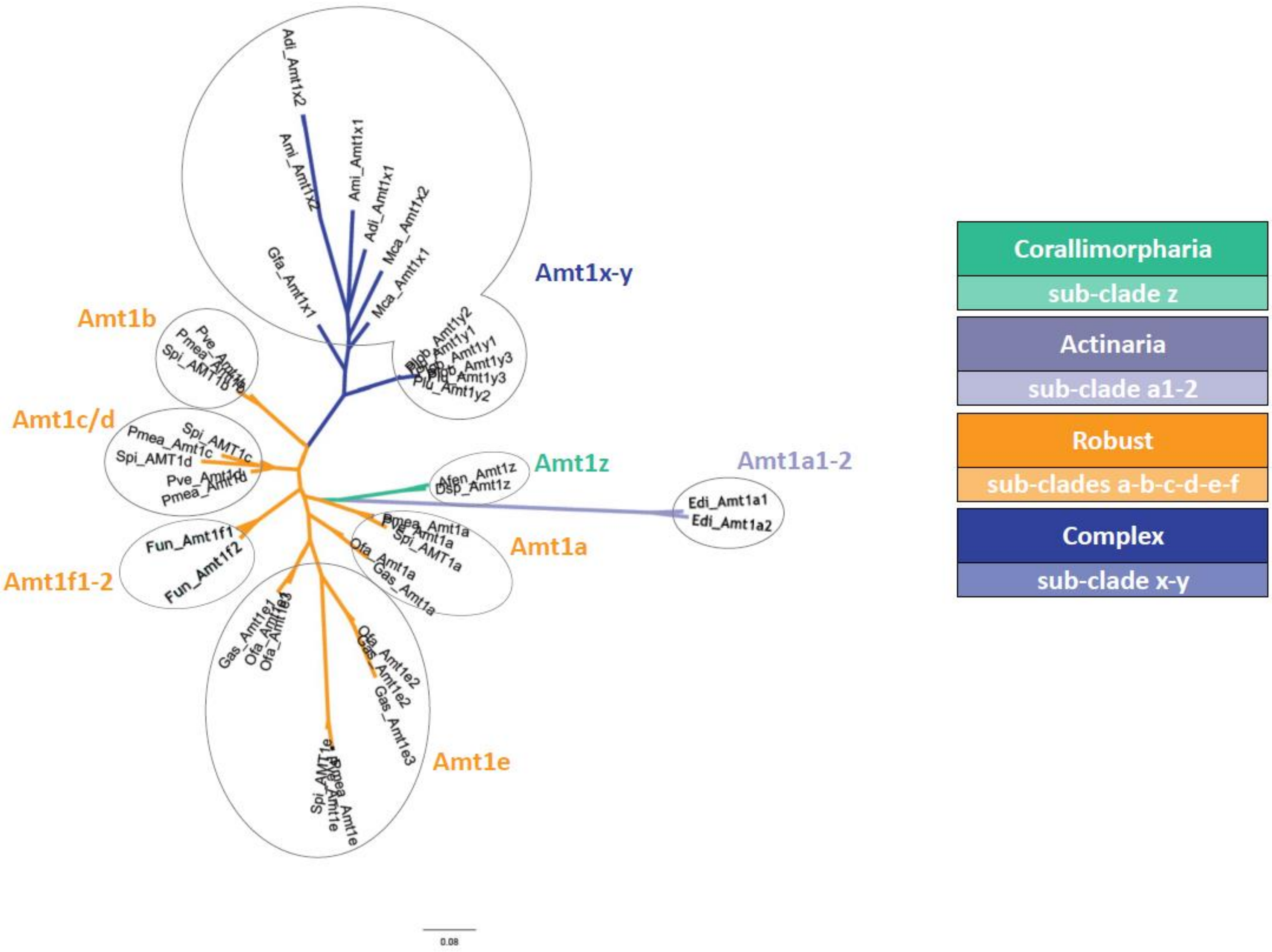

**Figure S30.** **Genomic localisation of cnidarian ammonium transporters Amt1 in various species**

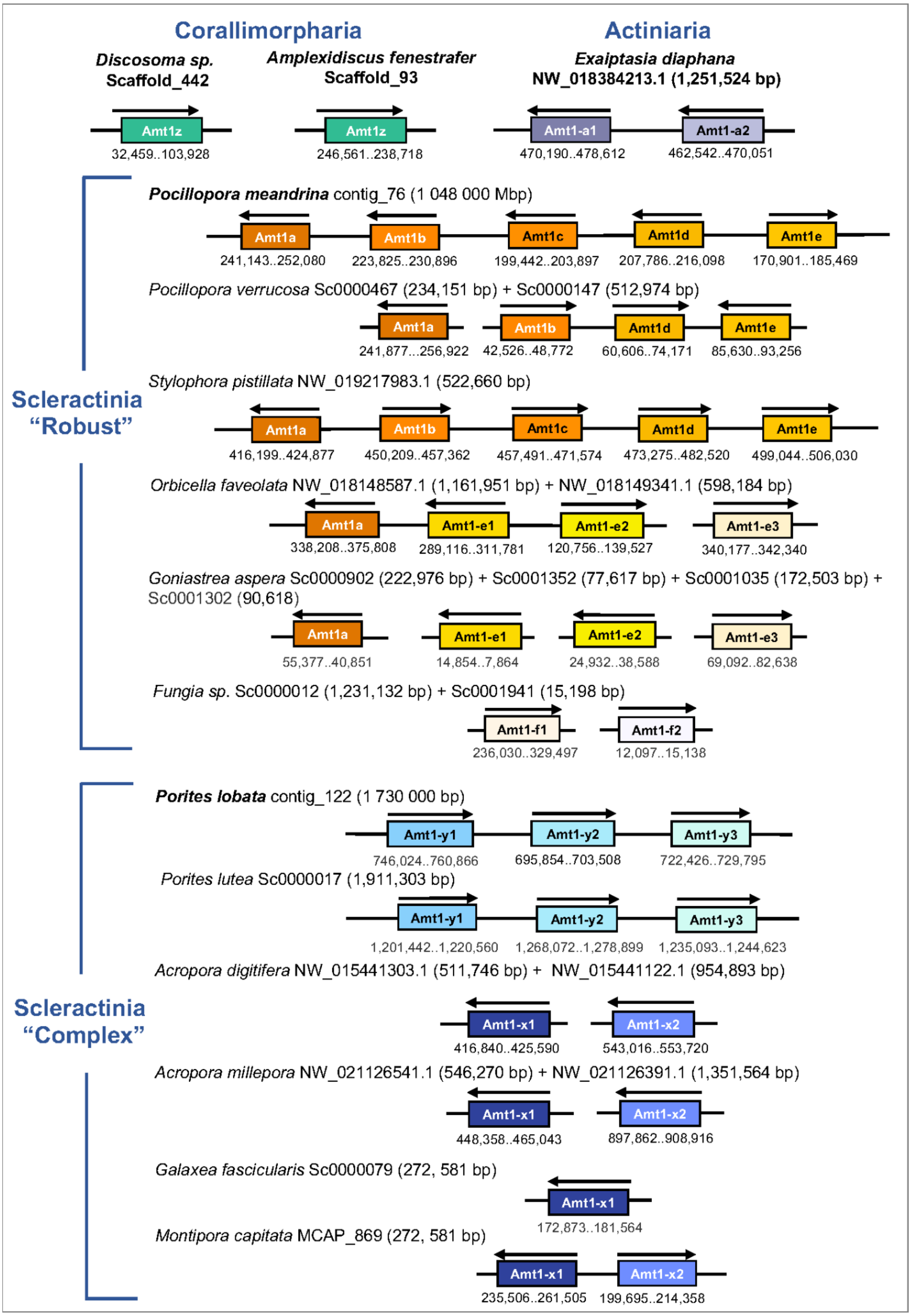

**Figure S31.** **Genomic localisation of Bicarbonate Anion Transporter SLC4 in various species.**

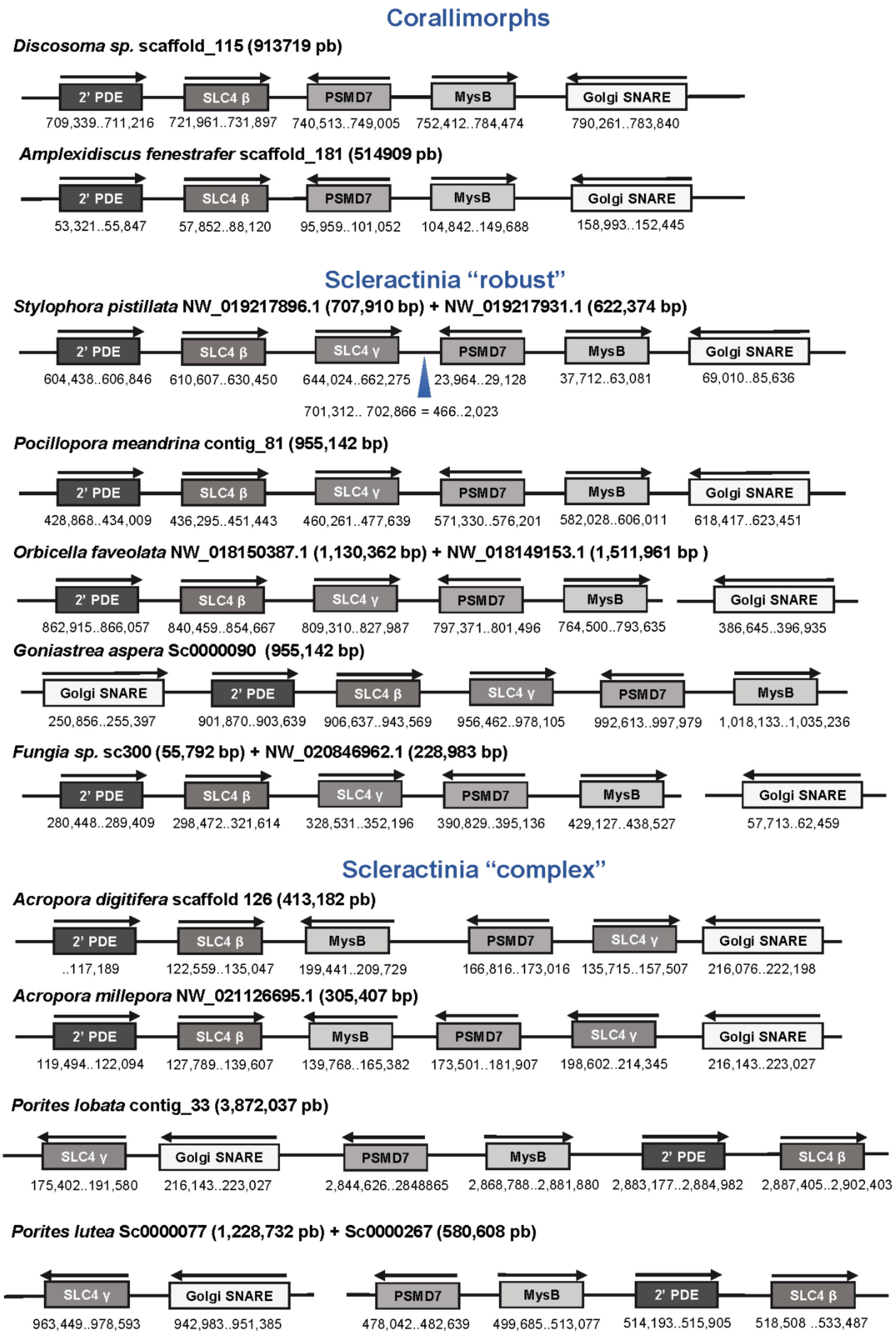

**Figure S32. Coral monocopy gene set.** Boxplots representing for each species the distribution of depth of mapping of short reads on 705 monocopy OG consensus (present in all coral species), normalized by the average depth on the 705 monocopy OGs consensus (A) or normalized by the average depth on 877 Busco metazoa ancestral sequences (B).

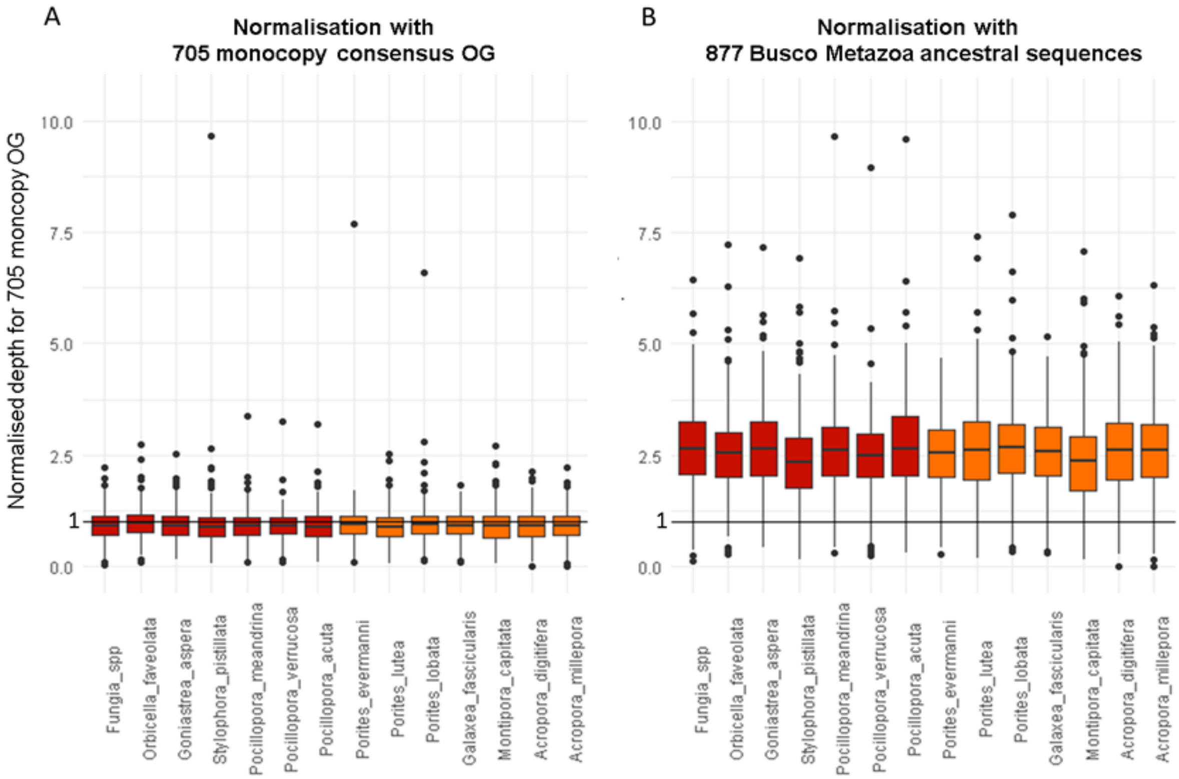
